# *Staphylococcus aureus* PSMα3 Cross-α Fibril Polymorphism and Determinants of Cytotoxicity

**DOI:** 10.1101/452011

**Authors:** Einav Tayeb-Fligelman, Nir Salinas, Orly Tabachnikov, Meytal Landau

**Affiliations:** Department of Biology, Technion-Israel Institute of Technology, Haifa 3200003, Israel.

**Author notes:** Departments of Chemistry and Biochemistry and Biological Chemistry, UCLA-DOE Institute, UCLA, Los Angeles, California 90095, USA.

**Keywords:** PSMα3, cytotoxins, cross-α fibrils, amyloid, membrane rupturing, co-aggregation, structural polymorphism, chameleon switch peptide.

## Abstract

The phenol-soluble modulin (PSM) peptide family, secreted by *Staphylococcus aureus,* performs various virulence activities, some mediated by the formation of amyloid fibrils of diverse architectures. Specifically, PSMα1 and PSMα4 structure the *S. aureus* biofilm by assembling into robust cross-β amyloid fibrils. PSMα3, the most cytotoxic member of the family, assembles into cross-α fibrils in which α-helices stack into tightly mated sheets, mimicking the cross-β architecture. Here we demonstrated that massive T-cell deformation and death is linked with PSMα3 aggregation and co-localization with cell membranes. Our extensive mutagenesis analyses supported the role of positive charges, and especially Lys17, in interactions with the membrane, and suggested their regulation by inter- and intra-helical electrostatic interactions within the cross-α fibril. We hypothesize that PSMα3 cytotoxicity is governed by the ability to form cross-α fibrils and involves a dynamic process of co-aggregation with cell membrane, rupturing it.

**Highlights:**

- The cytotoxic *S. aureus* PSMα3 assembles into cross-α fibrils
- Cross-α fibril polymorphism and mutations-induced secondary structure switching
- Regulation by cross-α fibril inter- and intra-helical electrostatic interactions
- Toxicity as a putative dynamic process of PSMα3 co-aggregation with membranes

## Introduction

Phenol-soluble modulins (PSMs) comprise a family of virulent peptides secreted by the pathogenic bacterium *Staphylococcus aureus* (Mehlin et al., 1999; Otto, 2014). PSMs stimulate inflammatory responses, contribute to biofilm formation and are capable of lysing human cells, including erythrocytes and leukocytes, such as T cells (Cheung, Joo, et al., 2014; Laabei et al., 2014). Several PSM members were shown to form amyloid fibrils, essential to their activity (Marinelli et al., 2016; Salinas et al., 2018; Schwartz et al., 2012; Tayeb-Fligelman et al., 2017). Amyloids are structured protein aggregates mostly known for their association with systemic and neurodegenerative diseases (Eisenberg and Jucker, 2012; Knowles et al., 2014; Tycko, 2015), but are also involved in various physiological functions in humans and in microbes (Chapman et al., 2002; DePas and Chapman, 2012; Fowler et al., 2005; Hughes et al., 2018; Jacob et al., 2016; Maji et al., 2009; Otzen and Nielsen, 2008; Pham et al., 2014; Schwartz et al., 2012; Soragni et al., 2015). They are known to form highly stable cross-β fibrils composed of tightly mated β-sheets, with β-strands oriented perpendicularly to the fibril axis (Eisenberg and Sawaya, 2017; Knowles et al., 2007). PSMα1 and PSMα4 comprise the robust cross-β fibrils, which confer high stability to the bacterial biofilm (Marinelli et al., 2016; Salinas et al., 2018; Schwartz et al., 2012), rendering it a resilient physical barrier against antibiotics and the immune system. However, PSMα3, the most cytotoxic member of the family (Cheung, Joo, et al., 2014; Cheung, Kretschmer, et al., 2014; Wang et al., 2007), forms unique cross-α fibrils (Tayeb-Fligelman et al., 2017). These fibrils are composed of amphipathic α-helices oriented perpendicularly to the fibril axis, which stack to form elongated ‘sheets’, stabilized by inter-helical polar interactions between side chains. The ‘helical sheets’ mate via an extensive hydrophobic core, providing stability to the fibril (Tayeb-Fligelman et al., 2017). PSMα3 cross-α fibrils exhibit unbranched fibril morphology, bind the amyloid dye-indicator thioflavin T (ThT), and are highly toxic to human T2 cells (Tayeb-Fligelman et al., 2017). Taken together, the secondary-structure polymorphism in amyloid-like structures formed by the homologous PSM family members (PSMα1, PSMα4 and PSMα3) is exceptional within dozens of amyloids studied thus far and is indicative of the specific function of each member.

The important role of PSMα3 in aggressive *S. aureus* infections (Wang et al., 2007), its unique cross-α fibril architecture and toxicity to human cells (Tayeb-Fligelman et al., 2017), encouraged us to further explore its structure-function-fibrillation relationships in order to identify the critical determinants in its mode of action. Using an array of derivatives and mutations, and assessment of their toxicity, structural and biophysical features, we herein demonstrate that the cross-α fibrillation process and positively charged residues are crucial for cytotoxicity. Specifically, we suggest that cross-α fibrillation regulates intra- and inter-helical electrostatic interactions to modulate activity, and that Lys17 is especially crucial for cytotoxicity, as it provides the only positive charge along the fibril available to interact with cell membrane lipid components. Moreover, we show that PSMα3 aggregates on the plasma membrane, and co-localizes with membrane particles of affected cells. The presented elucidation of structural and functional properties of the highly virulent PSMα3 is expected to provide a template for the development of novel antivirulence agents to combat *Staphylococcal* infections.

## Results

### Fibrillation and positive charges are key determinants of PSMα3 cytotoxicity

To identify the sequence and structural determinants crucial for PSMa3 fibrillation and activity, several alanine substitutions were generated and tested for their fibril-forming capacities (F3A, L7A, F8A, K9A, F10A, F11A, K12A, D13A, L15A, G16A, and K17A) (Table 1). Among these mutants, only F3A and F8A failed to form fibrils, at least as readily as the WT and other mutants, as visualized via transmission electron microscopy (TEM) (Figure 1A), demonstrating the robust tendency of PSMα3 to fibrillate. The non-fibrillating F3A and F8A mutants showed significantly reduced toxicity towards T2 cells (Figure 1B), while alanine substitutions in other non-polar residues (L7A, F10A, F11A, L15A, and G16A), along with D13A, maintained fibril-formation (Figure 1A) and toxic properties (Figure 1B). Moreover, introduction of a cysteine residue to either termini of PSMα3 showed that while C-cys-PSMα3 formed fibrils with fast kinetics and retained cytotoxicity, N-cys-PSMα3 lost the ability to fibrillate and correspondingly, was not cytotoxic (Figure S1). Furthermore, alanine substitutions of positively charged lysine residues (K9A, K12A, K17A) significantly reduced toxicity (Figure 1B), in accordance with a previous report (Cheung, Kretschmer, et al., 2014). These lysine to alanine mutants maintained the ability to form fibrils (Figure 1A), indicating the role of positively charged residues in cytotoxicity but not in fibrillation (Table 1), and that fibrillation alone is not sufficient.

**Figure 1.**
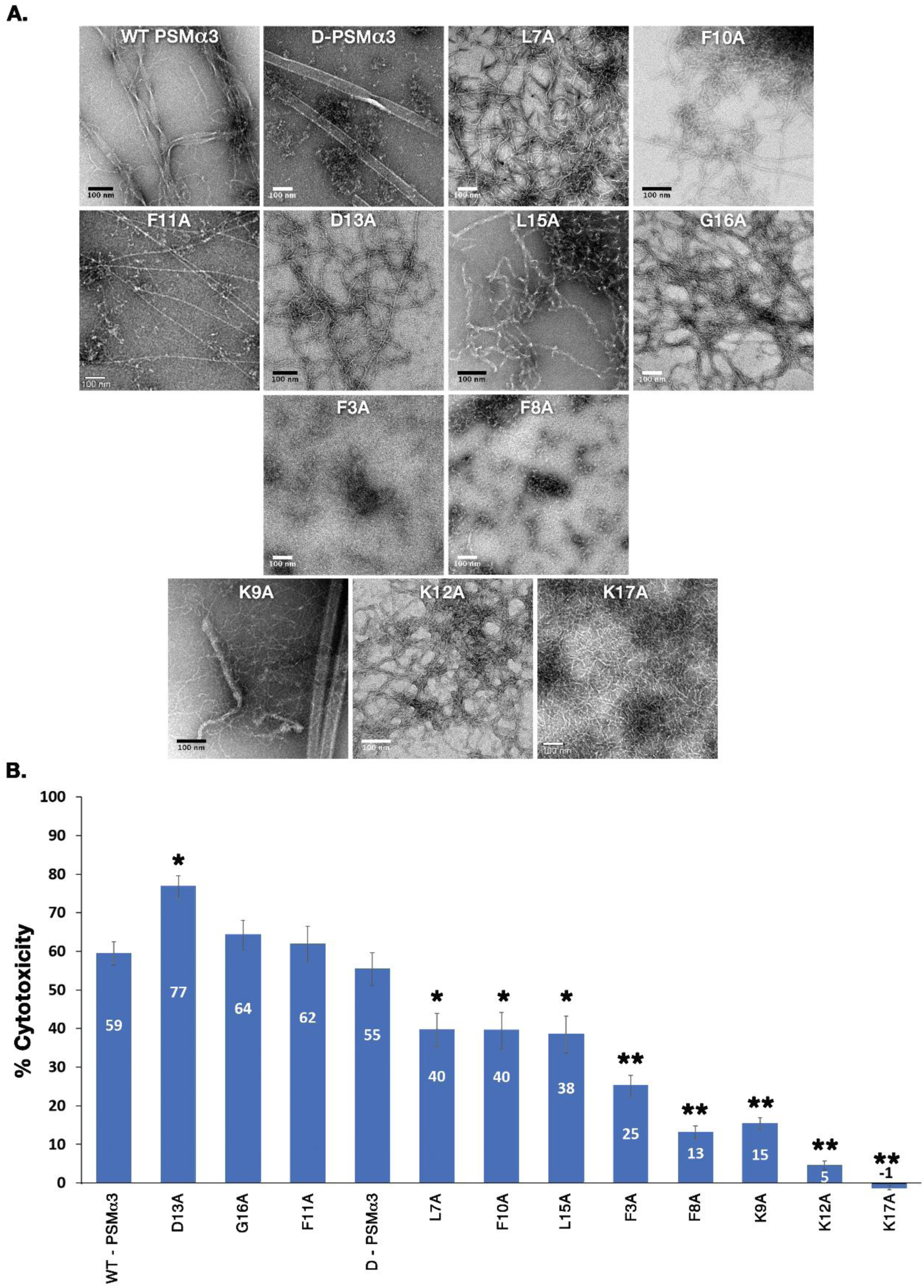
Fibril formation of PSMα3 derivatives and their toxicity against T2 cells. **A**. Fibril formation of PSMα3 mutants as visualized by transmission electron microscopy (TEM). WT PSMα3 and derivatives demonstrating the highest cytotoxicity levels, including D-PSMα3, L7A, F10A, F11A, D13A, L15A and G16A, form fibrils with various morphologies (top two rows). The F3A and F8A mutants (third row), showing diminished toxicity, did not form fibrils. The K9A, K12A and K17A mutants (bottom row), which also display diminished toxicity, formed fibrils but of reduced net positive charge. A scale bar of 100 nm is presented in all images. **B**. PSMα3-induced cell death was assessed using the lactate dehydrogenase (LDH) colorimetric assay upon 30 min of incubation of T cells with 4 µM of each PSMα3 variant. The experiment was repeated at least three times on different days, with similar results. The error bars represent the standard error of the mean. * p<0.05 and ** p<0.001 compared to WT PSMα3, accounting for the data from all three repeats.

**Table 1.** The biophysical and toxic properties PSMα3 and its derivatives.

| Peptide | Fibrils visualized by TEM** | ThT binding *** | Relative cytotoxicity ** (a) | Main secondary structure in solution (CD)*** | Secondary structure in the fibril (FTIR)*** | Main reflections in X-ray fiber diffraction** | Interpretation of fiber diffraction pattern |
| --- | --- | --- | --- | --- | --- | --- | --- |
| **WT PSMα3\*** | Fibrils | + | +++ | α-helical | α-helical | 10.5/12/25 Å | Cross-α |
| **D-PSMα3** | Fibrils | + | +++ | α-helical | N.D. | 10/12/25 Å | Cross-α |
| **F3A** | No fibrils | - | + | α-helical | N.D. | Diffuse | Ambiguous |
| **L7A** | Fibrils | - | ++ | α-helical | α-helical with partial transition to β-rich | 4.7/11.5/9-12/28.5 Å | A mixture of cross-α/$\beta$ |
| **F8A** | No fibrils | - | + | α-helical | N.D. | Diffuse | Ambiguous |
| **K9A** | Fibrils and crystals | + | + | α-helical with reduced helicity compared to WT | α-helical with partial transition to β-rich | 4.6/10/12/24 Å | A mixture of cross-α/$\beta$ |
| **F10A** | Fibrils | - | ++ | α-helical | N.D. | Diffuse | Ambiguous |
| **F11A** | Fibrils | - | +++ | α-helical with reduced helicity compared to WT | N.D. | 4.7/10/12.5/23-32 Å | A mixture with predominant cross-α diffraction |
| **K12A\*** | Fibrils | - | + | α-helical with reduced helicity compared to WT | α-helical with partial transition to β-rich | 4.3/9 Å | Could not interpret |
| **D13A** | Fibrils | - | ++++ | α-helical with reduced helicity compared to WT | α-helical with partial transition to β-rich | Diffuse | Ambiguous |
| **L15A\*** | Fibrils | + | ++ | α-helical | N.D. | 4.6/10.5/12.5/25 Å | A mixture with predominant cross-α diffraction |
| **G16A\*** | Fibrils | + | +++ | α-helical | N.D. | Diffuse | Ambiguous |
| **K17A** | Fibrils | + | - | α-helical with reduced helicity compared to WT | α-helical | 10/12.5/25 Å | Cross-α |
| **C-cys-PSMα3** | Fibrils | + | +++<sup>(b)</sup> | N.D. | N.D. | N.D. | N.D. |
| **N-cys-PSMα3** | No fibrils | - | +<sup>(b)</sup> | N.D. | N.D. | N.D. | N.D. |

N.D. - Not determined
\* Crystal structure solved and showed cross-α fibrils
\*\* No pre-treatment of the peptides
\*\*\* Peptides were pretreated to dissolve preformed aggregates, as detailed in the Methods section
(a) Results not influenced by peptide pretreatment.
(b) Toxicity induced by 8 μM peptide; all other peptides were tested at 4 μM

WT PSMα3, and the K9A, L15A, G16A, and K17A mutants displayed various fibrillation kinetics, as assessed by monitoring ThT fluorescence (Figure S2). However, other mutants (L7A, F10A, F11A, K12A and D13A) failed to bind ThT (Figure S2), as was previously reported (Zheng et al., 2018), despite clear fibril-forming ability visualized by TEM (Figure 1A). These findings emphasize the importance of integrating different methods to assess fibrillation propensity, and accordingly, point to fibrillation and positive charges as the dominant factors governing cytotoxicity.

### Secondary structure properties of PSMα3 mutants in soluble and fibrillar states

The secondary structure properties of PSMα3 mutants were explored in the soluble state and in fibrils. WT PSMα3, as well as most mutants substituting non-polar residues (F3A, L7A, F10A, L15A and G16A), and the all D-enantiomeric amino acid PSMα3 (D-PSMα3), maintained helicity in solution, measured using circular dichroism (CD) (Figure 2 and Figure S3). In contrast, mutants in which a polar residue was substituted to alanine (K9A, K12A, K17A, and D13A), thereby disturbing the amphipathic nature of the peptide, exhibited reduced helicity in solution compared to the WT PSMα3 (Figure 2).

**Figure 2.**
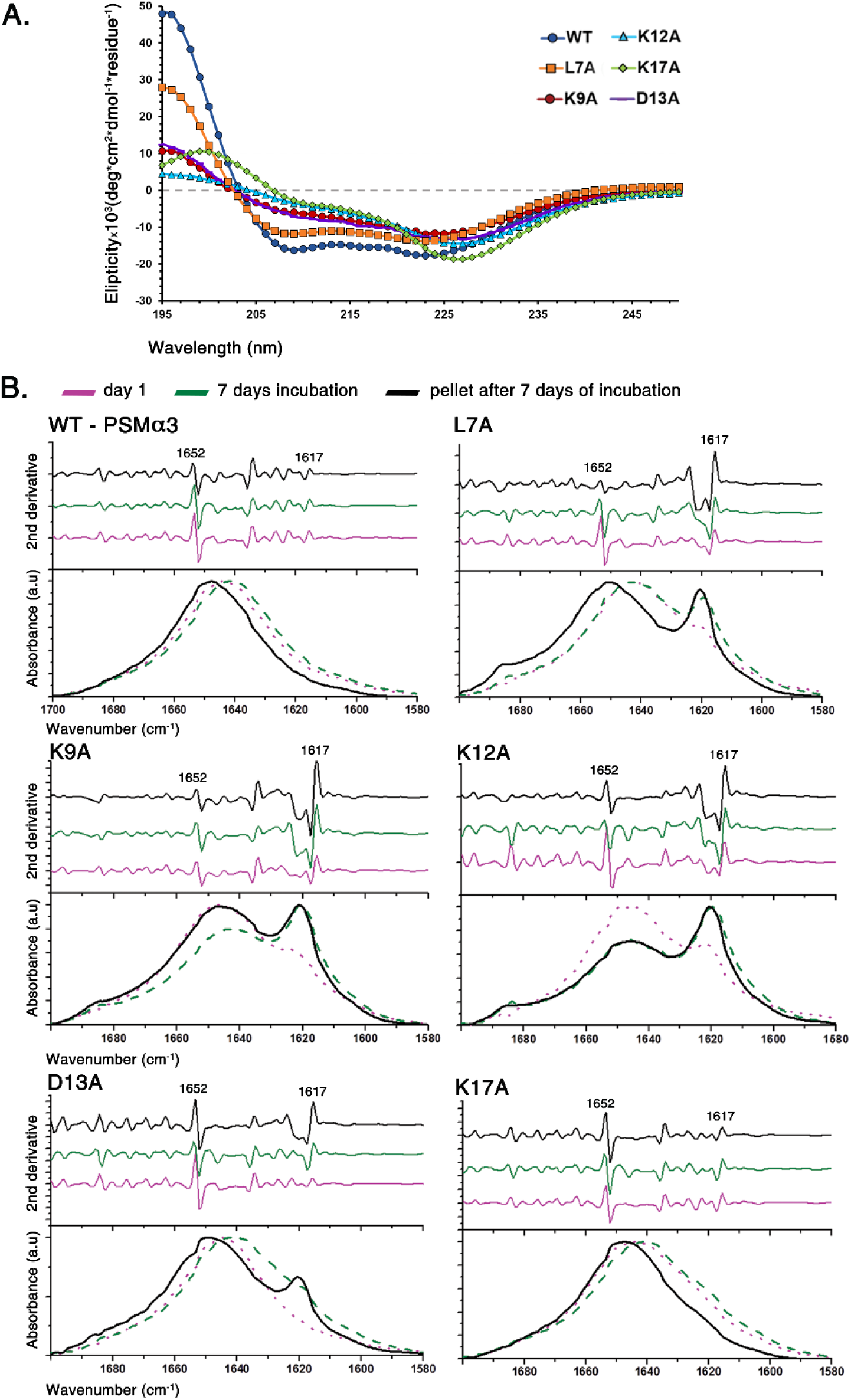
Secondary structure analyses of WT PSMα3 and mutants with polar residue substitutions. **A**. The CD spectrum of WT PSMα3 shows helical configuration with minima at 208 and 222 nm. The K9A, K12A, D13A and K17A mutants demonstrate diminished helicity compared to WT PSMα3. The measurements were done using 111 µM peptides in a buffer containing 150 mM sodium chloride and 10 mM sodium phosphate at pH 7.4. The concentrations of the peptides were calculated during the measurement to account for the effect of aggregation, as described in the Experimental section. **B**. ATR-FTIR spectroscopy analyses of the peptides, dissolved at 25 mg/ml (∼10 mM), upon solubilization (day one, dotted pink line), after one week of incubation at room temperature (day seven, dashed green line), and isolation of the pellet from the seven-day incubation (pellet, smooth black line). The experiments were repeated three times and the amide I’ region of representative curves is shown. The secondary derivative of each curve was calculated and depicted to resolve and identify the major peaks contributing to the ATR-FTIR overlapping signal in this region. Peaks in the region of 1645–1662 cm^-1^ indicate the existence of α-helices, 1637–1645 cm^-1^ indicate disordered species, 1630–1643 cm^-1^ correlate with small and disordered β-rich amyloid fibrils with absorbance which is typical of bent β-sheets in native proteins, and 1611–1630 cm^-1^ as well as ∼1685–1695 cm^-1^ are indicative of rigid cross-β fibrils (Goormaghtigh et al., 1990; Moran and Zanni, 2014; Sarroukh et al., 2013; Zandomeneghi et al., 2009).

The secondary structures of the WT PSMα3 and its K17A mutant in solid state, analyzed using ATR-FTIR (Figure 2), showed a main peak at 1652 cm^-1^, indicating on a predominant α-helical content (Goormaghtigh et al., 1990; Moran and Zanni, 2014; Sarroukh et al., 2013; Zandomeneghi et al., 2009), when dried as a thin film on the ATR module, both on the day of solubilization, and after seven days of incubation in solution. Helicity was even more dominant in isolated fibrils obtained by washing the fibril pellets after seven days of incubation (Figure 2). In contrast, the L7A, K9A, K12A and D13A mutants showed some β-rich content upon incubation, especially in the fibril pellet, with an additional FTIR peak at 1617 cm^-1^ (Figure 2), indicating on rigid cross-β fibrils (Goormaghtigh et al., 1990; Moran and Zanni, 2014; Sarroukh et al., 2013; Zandomeneghi et al., 2009).

The X-ray fiber diffraction of WT PSMα3 and of D-PSMα3, displayed the cross-α diffraction pattern, with orthogonal reflection arcs at ∼10.5 Å and ∼12 Å (Figure 3), corresponding to the spacing between α-helices along each sheet, and the inter-sheet distance, respectively, in the PSMα3 crystal structure (Figure 4). An additional major reflection was present at ∼25 Å, which may be attributable to the length of the α-helix, or to the inter-sheet distance over the dry and wet interfaces (Table S1). The X-ray fiber diffraction profile of the K17A mutant showed a very similar cross-α pattern (Figure 3), in accordance with the helicity observed for WT PSMα3 and K17A fibrils in the FTIR spectra (Figure 2). The diffraction pattern of the L7A, K9A, F11A, and L15A mutants indicated additional structural features characteristic of cross-β structures, with reflections at 4.6-4.7 Å and 9-12 Å (Figure 3). This was especially pronounced in the moderately toxic L7A mutant which showed additional strong orthogonal reflections at 4.7 Å and 9-12 Å (Figure 3). The diffraction pattern of these four mutants may be indicative of mixed populations of both cross-α and cross-β (Table 1). Other mutants showed only diffuse reflections that could not be interpreted (Figure S4). Taken together, WT PSMα3 and its K17A mutant consistently maintained α-helical conformation in solution and in fibrils, whereas the fibrils of the L7A, K9A, F11A, K12A, D13A and L15A mutants demonstrated both helical and β-rich features suggesting a mixed population (Table 1).

**Figure 3.**
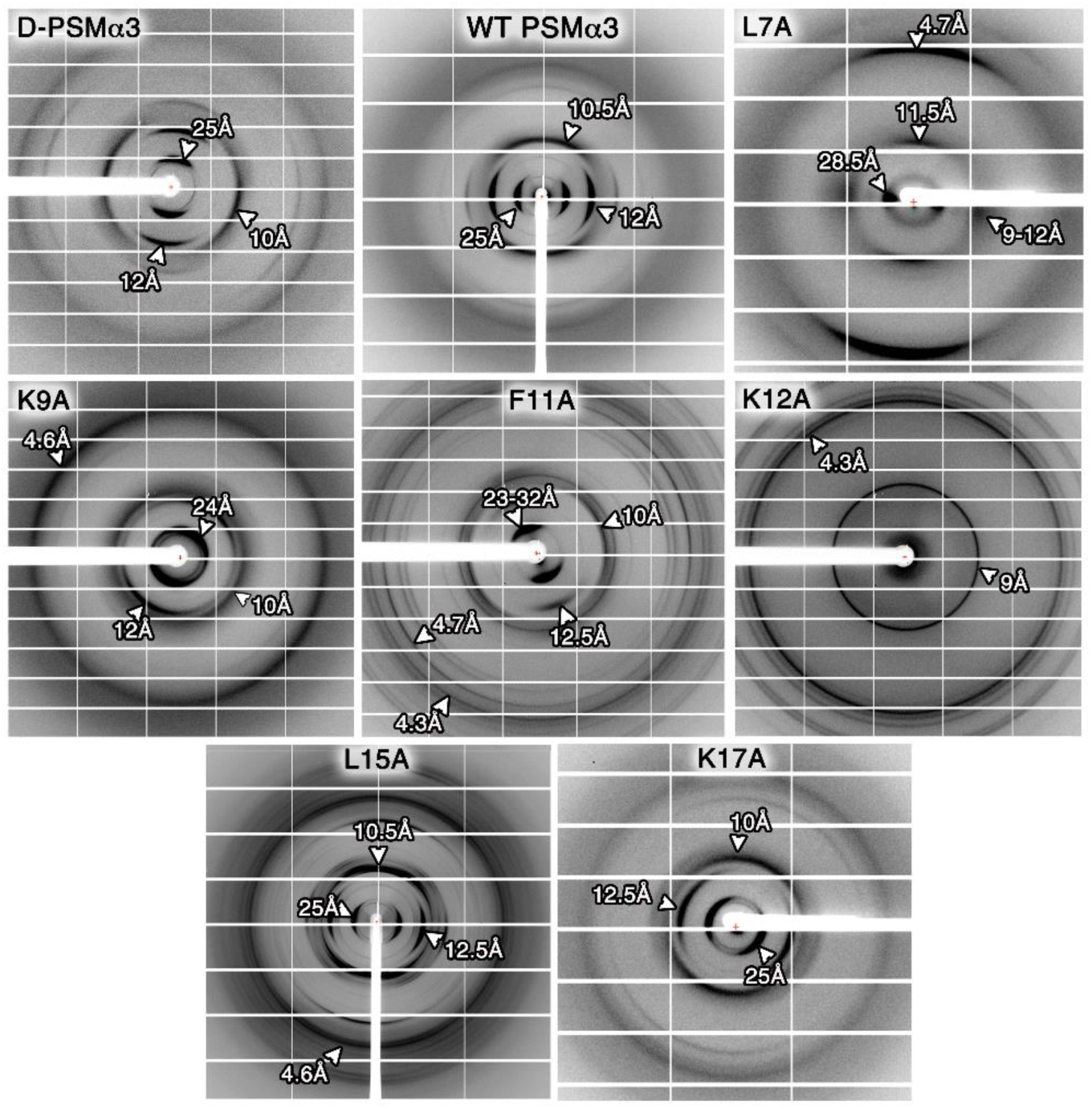
X-ray fiber diffraction of PSMα3 and derivatives. X-ray fiber diffraction patterns, with major reflections labeled. Peptide powders were dissolved in ultra-pure water to a concentration of 20 mg/ml and a microliter drop was applied between two sealed-end glass capillaries. The fiber diffraction was measured upon drying of the fibrils or after up to 10 hours of incubation on the capillaries.

**Figure 4.**
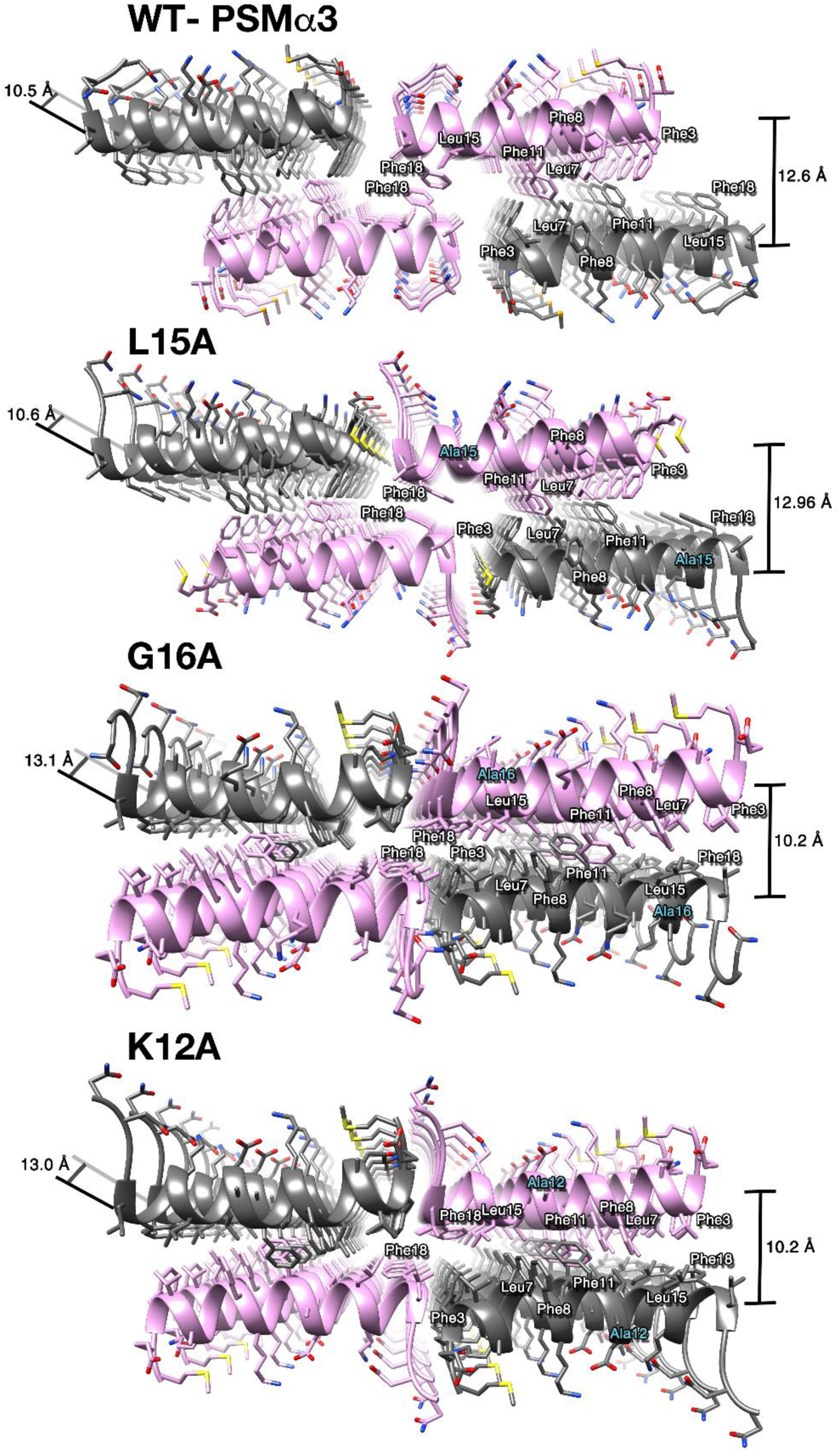
The crystal structures of PSMα3 and its mutants display packing polymorphism. The crystal structures of PSMα3 (PDB ID 5I55) (Tayeb-Fligelman et al., 2017), and the L15A (PDB ID 6GQ5), G16A (PDB ID 6GQC) and K12A (PDB ID 6GQ2) mutants determined at 1.45 Å, 1.5 Å, 1.4 Å and 1.5 Å resolution, respectively. The view is down the fibril axis. Two pairs of sheets are depicted in gray or orchid ribbons, side chains are represented as sticks and labeled by their three-letter amino acid code and position number. Heteroatoms are colored by atom type, and alanine substitution sites are indicated by cyan labels. The mated sheets form a hydrophobic dry interface (lacking water molecules) with inter-sheet distances shown on the right. The distances between α-helices along each sheet are shown on the left.

### Crystal structures of three PSMα3 mutants reveal packing polymorphism in their cross-α architecture

The crystal structure of the K12A, G16A, and L15A PSMα3 mutants demonstrated cross-α fibrils of tightly mated sheets composed of parallel amphipathic α-helices facing each other via their hydrophobic face (Figure 4, Table 2). The structures demonstrated packing polymorphism, namely translations of the mated sheets from a staggered orientation of slightly shifted sheets in the WT PSMα3 and the L15A mutant, to a directly facing orientation in the K12A and G16A mutants. The structures of WT PSMα3 and L15A also displayed a larger inter-sheet distance and a smaller distance between the α-helices along the sheets, in comparison to G16A and K12A (Figure 4 and Table S1).

**Table 2.** Data collection and refinement statistics.

| | PSMα3 L15A | PSMα3 K12A | PSMα3 G16A |
| --- | --- | --- | --- |
| Beamline | ESRF ID30a-3 | ESRF ID30a-3 | ESRF ID23_2 |
| Date | November 10<sup>th</sup> 2016 | November 10<sup>th</sup> 2016 | September 17<sup>th</sup> 2016 |
| PDB ID | 6GQ5 | 6GQ2 | 6GQC |
| **Data collection** |  |  |  |
| Space group | P1 21 1 | C 1 2 1 | C 1 2 1 |
| **Cell dimensions** |  |  |  |
| a, b, c (Å) | 28.41 10.67 29.60 | 52.94 13.15 25.97 | 53.27 13.30 25.80 |
| $\alpha$, $\beta$, $\gamma$ (°) | 90.0 105.9 90.0 | 90.0 112.3 90.0 | 90.0 111.9 90.0 |
| Wavelength (Å) | 0.96770 | 0.96770 | 0.8729 |
| Resolution (Å) | 28.469-1.500 (1.54-1.50) | 24.496-1.540 (1.58-1.54) | 24.711-1.400 (1.44-1.40) |
| R-factor observed (%) | 13.7 (55.9) | 11.4 (83.6) | 12.4 (75.4) |
| <sup>a</sup>R<sub>meas</sub> (%) | 17.1 (69.2) | 14.2 (103.2) | 13.5 (82.2) |
| *I* / $\sigma$ *I* | 4.09 (1.56) | 4.83 (1.20) | 8.13 (2.28) |
| Total reflections | 7650 (549) | 6862 (468) | 21920 (1637) |
| Unique reflections | 2854 (207) | 2477 (172) | 3478 (267) |
| Completeness (%) | 96.3 (95.0) | 94.1 (91.5) | 99.0 (100.8) |
| Redundancy | 2.68 (2.65) | 2.77 (2.7) | 6.30 (6.13) |
| <sup>b</sup>CC1/2 (%) | 98.7 (85.1) | 98.3 (77.2) | 99.4 (84.9) |
| **Refinement** |  |  |  |
| Resolution (Å) | 17.47-1.50 (1.68-1.50) | 14.61-1.54 (1.72-1.54) | 14.67-1.40 (1.44-1.40) |
| Completeness (%) | 96.45 (97.42) | 94.58 (96.23) | 99.29 (100.00) |
| <sup>c</sup> No. reflections | 2568 (713) | 2229 (620) | 3129 (238) |
| <sup>d</sup>R<sub>work</sub> (%) | 19.4 (33.4) | 20.3 (37.9) | 15.1 (23.9) |
| R<sub>free</sub> (%) | 24.50 (36.1) | 23.6 (38.0) | 19.3 (26.4) |
| No. atoms | 201 | 189 | 199 |
| Protein | 182 | 181 | 194 |
| Ligand/ion | 8 | 5 | 3 |
| Water | 11 | 3 | 2 |
| **B-factors** |  |  |  |
| Protein | 17.428 | 25.961 | 17.827 |
| Ligand/ion | 41.255 (MRD) | 85.652 (SO<sub>4</sub>) | 43.355 (Na<sup>+</sup>)<br>53.659(Cl<sup>-</sup>) |
| Water | 23.327 | 33.584 | 24.645 |
| **R.m.s. deviations** |  |  |  |
| Bond lengths (Å) | 0.019 | 0.013 | 0.011 |
| Bond angles (°) | 1.692 | 1.419 | 1.385 |
| Clash score (Chen et al., 2010) | 2.55 | 0.00 | 0.00 |
| Molprobit score (Chen et al., 2010) | 1.04 | 0.5 | 1.03 |
| Molprobit percentile (Chen et al., 2010) | 99th percentile | 100th percentile | 99th percentile |
| Number of xtals used for scaling | One crystal | One crystal | One crystal |

Values in parentheses are for highest-resolution shell.
(a) R-meas is a redundancy-independent R-factor defined in Reference (Weiss and IUCr, 2001).
(b) CC<sub>1/2</sub> is the percentage of correlation between intensities from random half datasets (Karplus and Diederichs, 2012).
(c) Number of reflections corresponds to the working set.
(d) Rwork corresponds to the working set

### Electrostatic interactions within the cross-α fibril putatively regulate PSMα3 toxicity

The crystal structures of PSMα3 and its mutants provided the opportunity to explore properties stabilizing the α-helices and their interactions within the cross-α fibrils. Generally, the formation of intra-helical salt-bridges between positively and negatively charged residues located 3-4 positions apart on the sequence are associated with increased α-helical stability (Kumar and Nussinov, 2002; Laabei et al., 2014). In the twisted helical configuration of monomeric PSMα3 observed by NMR in solution (Towle et al., 2016), such an intra-helical salt bridge can form between Asp13 and Lys17. However, in the fibrillary crystal structures of PSMα3 and its mutants, demonstrating canonical α-helices, such salt bridge can form between Asp13 and Lys9. In the L15A mutant, an additional intra-helical electrostatic bond formed between Glu2 and Lys6. In PSMα3 and the L15A mutant fibrils, Lys12, from an adjacent α-helix in the same sheet, constitutes a possible second salt-bridge partner for Asp13, this time inter-helical, which may stabilize the fibril (Figure 5). In the structures of the WT and of the K12A and G16A mutants, an inter-helix salt bridge may form along the fibril between Glu2 and Lys6. In all four crystal structures, Lys17 is the only lysine residue fully deprived of potential salt-bridges (Figure 5). Accordingly, the K17A mutation, the least toxic peptide among all tested alanine mutations (Figure 1B), did not affect the electrostatic interactions within the fibril, leading to similar stability of its α-helical configuration in fibrils to that of WT PSMα3 (Figures 2&3). We therefore hypothesized that in the aggregated form, Lys17 contributes an exposed and available positive charge that might mediate cytotoxicity via membrane interactions. The D13A mutant eliminates potential electrostatic interactions with lysine residues and possesses an increased net positive charge, rendering it the most cytotoxic derivative (Figure 1B). Overall, our results provide further support of the proposed role of positively charged residues in cell membrane interactions and suggest that fibrillation regulates the availability of these charges to interact with the membrane.

**Figure 5.**
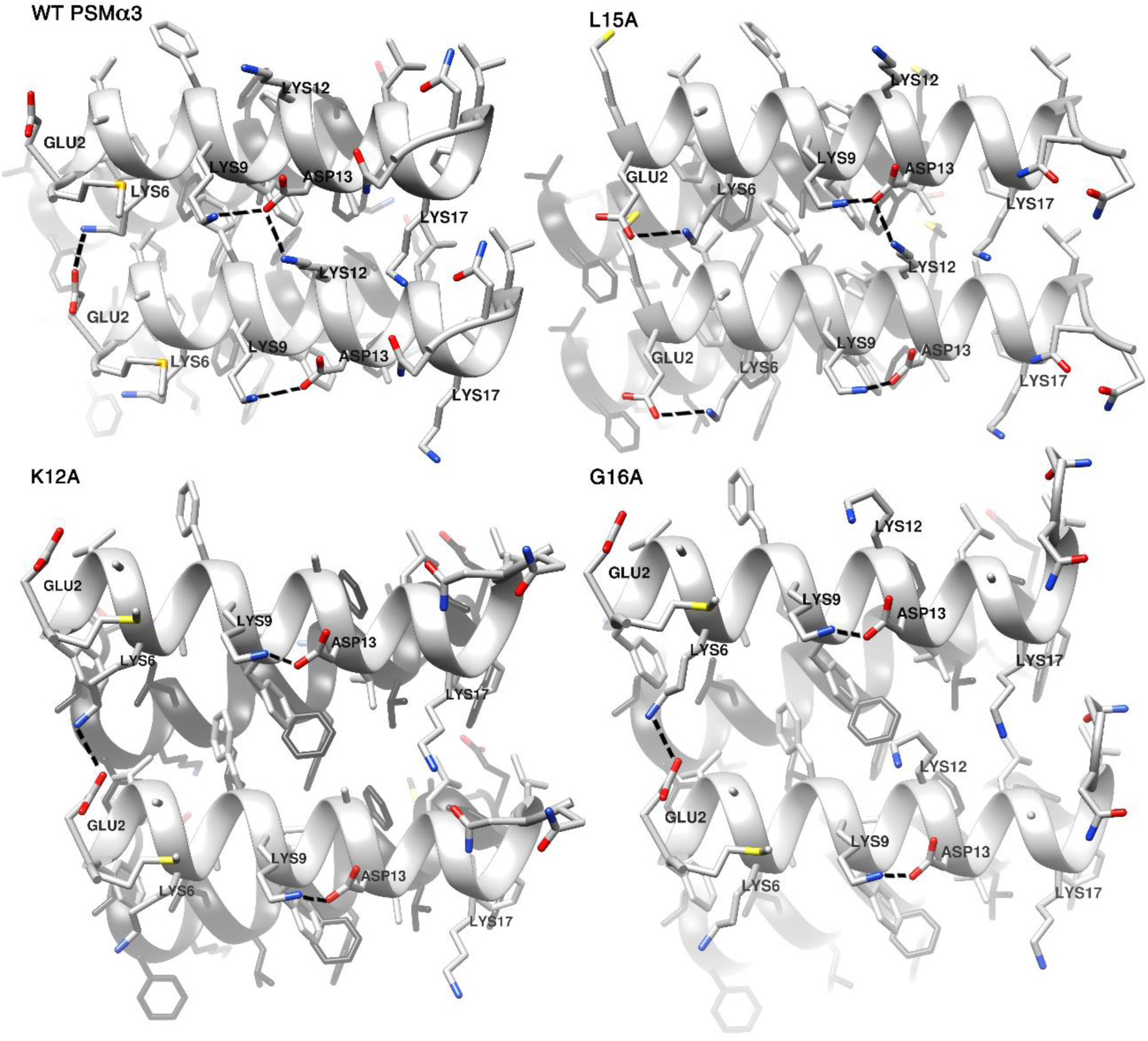
Potential salt bridges in the crystal structures of PSMα3 and its mutants. Crystal structures of PSMα3 and the K12A, L15A, and G16A mutants emphasizing potential intra- and inter-helical electrostatic interactions, with a view perpendicular to the fibril axis. Side chains are shown as sticks and labeled by their three-letter amino acid code and position number, and heteroatoms are colored according to atom type. Potential salt bridges are depicted by dashed lines.

### Aggregates of PSMα3 co-localize with cell membranes leading to cell death

Incubation of PSMα3 with T2 cells elicited PSMα3 aggregation on the cell membranes, as visualized by confocal microscopy using a mixture of WT PSMα3 and a fibrillating fluorescein isothiocyanate (FITC)-labeled PSMα3 derivative (Figure 6 & S5). Scanning electron micrographs demonstrated the massive deformation and rapid cell death occurring within a few minutes of exposure to PSMα3 (Figure 7). The interactions of PSMα3 with the membrane, and resulting membrane rupture and cell death, were also evident by the appearance of membrane particles co-localized with PSMα3 aggregates around affected cells, as observed by continuous live cell confocal microscopy (Figure 8). The process of co-localization and possible co-aggregation with the cell membrane are governed by the peptide’s ability to fibrillate, however requires the existence of soluble species to enable a dynamic process. Correspondingly, incubated PSMα3 shows an active process of fibrillation in a solution separated from fibrils following incubation (Figure S6), suggesting equilibrium between monomers, or soluble aggregates, and fibrils.

**Figure 6.**
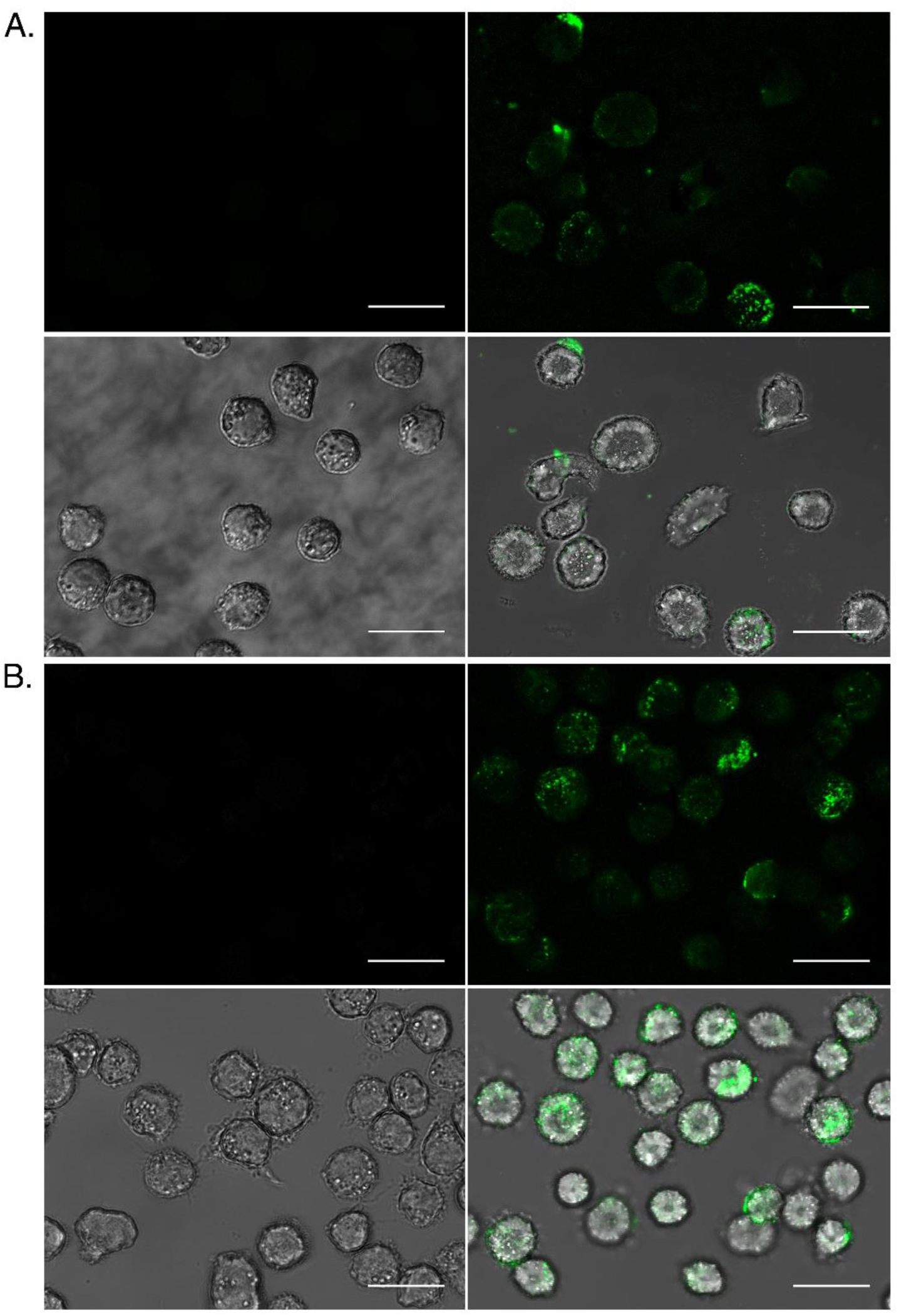
Confocal microscopy of PSMα3 aggregation on the cell membrane. Representative confocal microscopy images of untreated T2 cells (left panels), and cells incubated for 10 min (**A**) or 30 min (**B**) with a peptide mixture of 5 µM WT PSMα3 and 5 µM FITC-labeled PSMα3 (right panels). The FITC channel (top) was merged with the bright-field channel (bottom) to indicate peptide location in the cell. A 20 µm scale bar is shown for all images.

**Figure 7.**
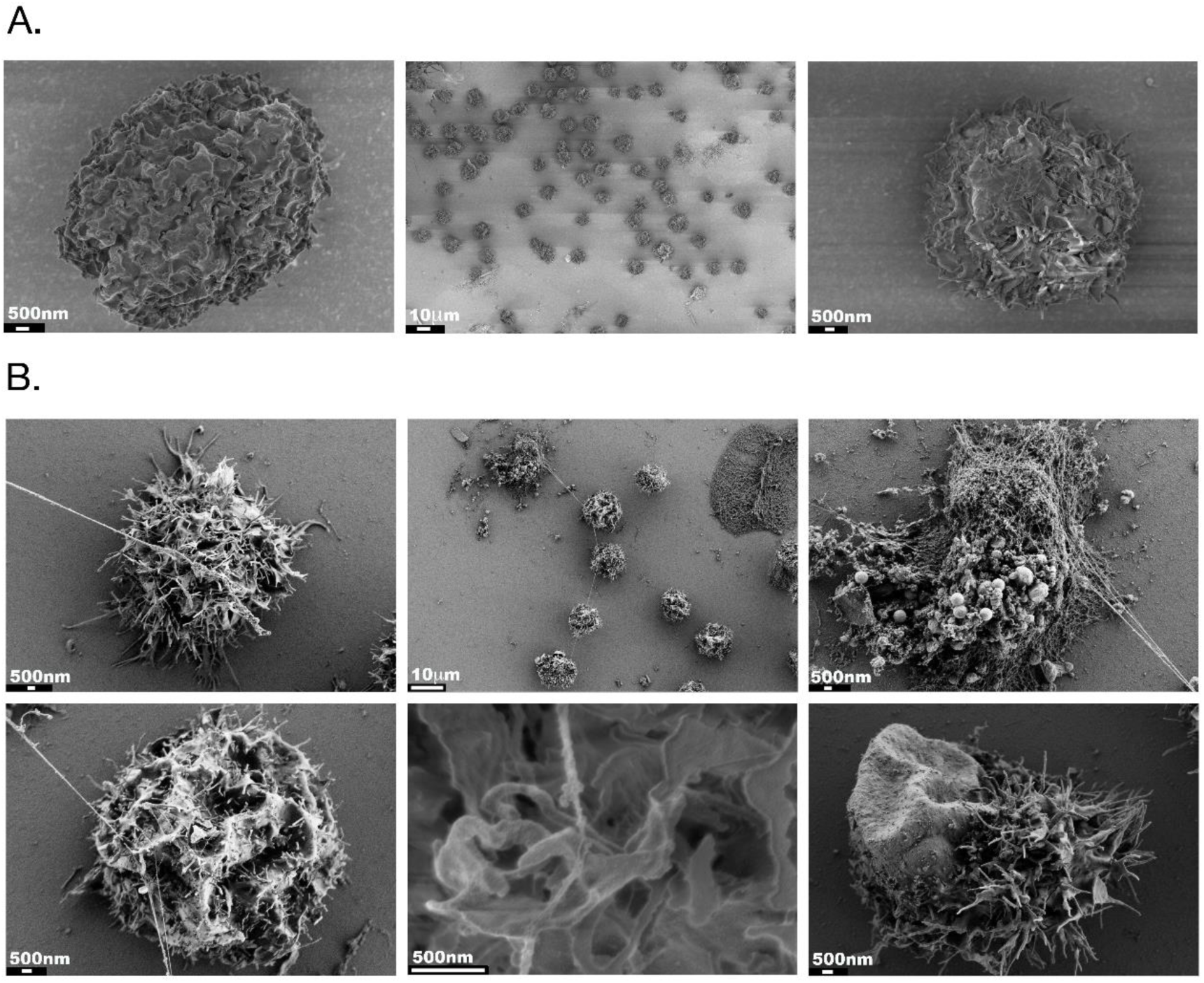
Scanning electron micrographs showing deformation of PSMα3-treated T2 cells. Scanning electron microscopy (SEM) images of (**Α**) untreated T2 cells, and (**B**) T2 cells incubated for 5 min with 10 µM PSMα3. The micrographs were taken at different magnifications, as indicated by the scale bars (white on a black background) (10 µm (zoom-out) or 500 nm (zoom-in)).

**Figure 8.**
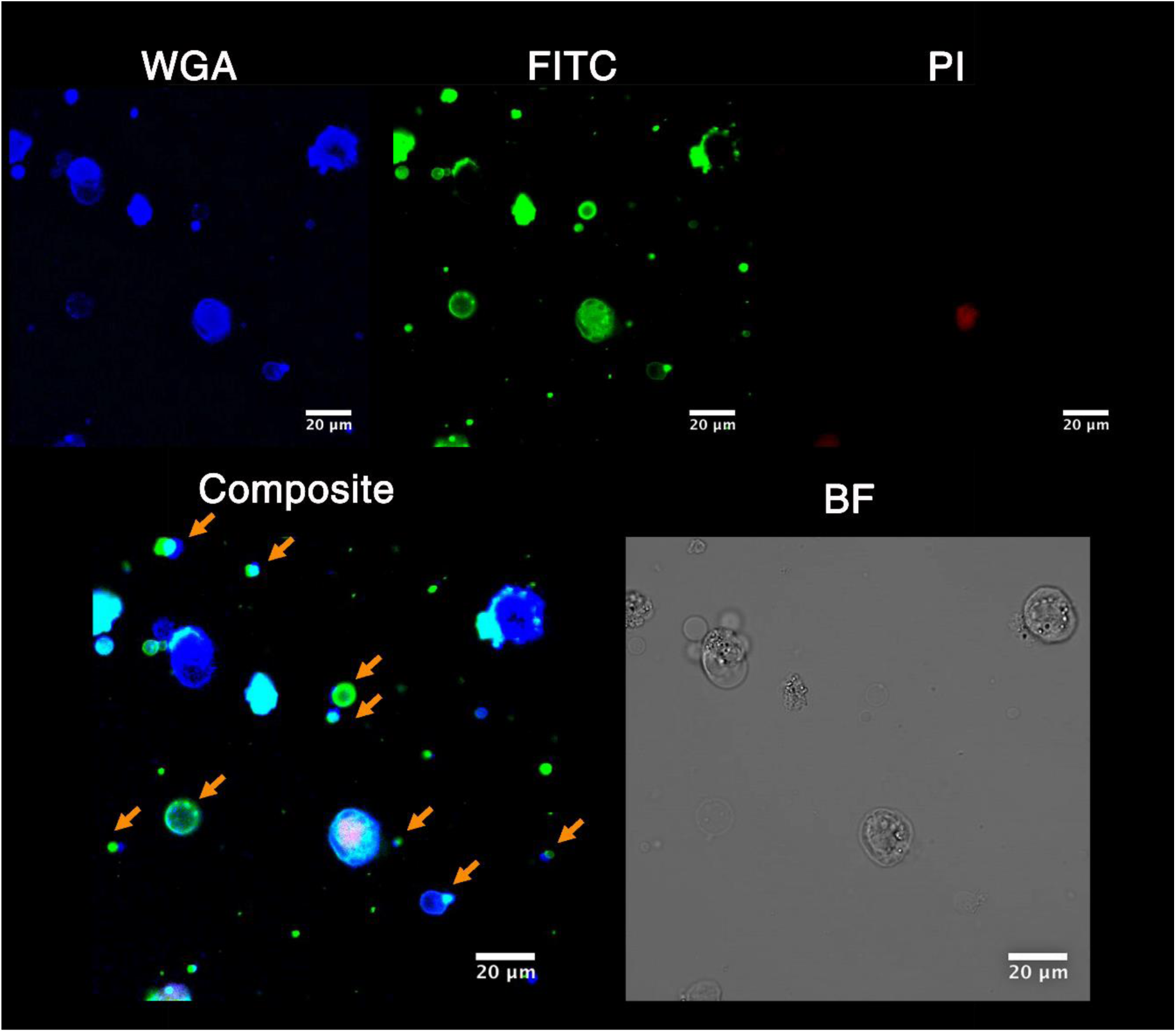
PSMα3 co-aggregates with membrane particles around affected cells. T2 cells were treated with a mixture of 5 µM WT and 10 µM FITC-labeled PSMα3 peptides (green) in the presence of propidium iodide (PI, red). The cell membranes were stained with wheat gram agglutinin (WGA, blue). Representative images from the live-cell imaging were taken after 50 min. In the composite image of the stains, orange arrows indicate particles of extracted cell membrane co-localized with the aggregated peptide. Bright field (BF) image show cell damage.

## Discussion

PSMα3, a cytotoxic peptide secreted by *S. aureus*, is acting as a functional cross-α fibril (Tayeb-Fligelman et al., 2017). The distinctive secondary structure observed for the PSMα3 cross-α fibrils, particularly when compared to PSMα1 and PSMα4, which are involved in *S. aureus* biofilm structuring, and transition to form the canonical cross-β amyloid fibrils (Salinas et al., 2018), is indicative of structurally-encoded functional specificity. The role of fibrillation in PSMα3 activities is yet not fully understood, and its impact on cytotoxicity, in particular, has been under debate (Yao et al., 2019; Zheng et al., 2018). The results presented here demonstrate a critical role of the fibrillation process in cytotoxicity and provide structural and mechanistic insight into the involvement of inter- and intra-helical polar interactions within the peptide assembly in regulating toxicity.

Fibrillation is a robust feature of PSMα3, as shown by preserved fibrillation capacities in nine of the eleven single point alanine mutations tested (Figure 1A). F3A and F8A, the two mutants that failed to form fibrils under our experimental conditions (or not as readily as the WT and other mutants, as these methods can never completely exclude fibril formation), were also the least toxic, with the exception of mutations in the lysine residue that provide another important determinant of toxicity. In addition, while the C-cys-PSMα3 derivative formed fibrils and was cytotoxic, the N-cys-PSMα3 did not form fibrils and was not cytotoxic (Figure S1). This radical difference between peptides with the same amino-acid composition endorses the central role of fibrillation in PSMα3 toxic activity. Furthermore, the significant effect of F3A, F8A and N-cys-PSMα3 on fibrillation and toxicity, and the structural changes observed for fibrils of the moderately toxic L7A mutant (Figure 3), may point to the N-terminal part as a critical region in cross-α fibrillation. The significantly reduced toxicity of the mutants substituting lysine residues (K9A, K12A and K17A), which retained fibril formation, suggest that fibrillation alone in not sufficient for toxicity. PSMα3 toxicity requires specific positive charges and perhaps also specific fibril properties, which were altered in the K9A and K12A mutants compared to the WT according to the secondary structure and fiber diffraction analyses (Figures 2-3). Other determinants of toxicity are still left to be determined, and although the results point to aggregation as promoting toxicity, it is yet impossible to determine the exact morphologies that interact with the cells. Moreover, since the effect of mutagenesis on toxicity varies with cell types (Cheung, Kretschmer, et al., 2014), differences in toxicity mechanisms can also be expected.

Polymorphism within cross-β fibrils is a known phenomenon of amyloids (Tycko, 2015), contrasting the dogma within globular proteins that structure is more conserved than sequence. We found polymorphism also in cross-α fibrils, observed between crystal structures of PSMα3 and the three single-point alanine mutants, L15A, G16A and K12A (Figure 4). While PSMα3 and the K17A mutant are strictly helical in fibrils, the L7A, K9A, F11A, K12A, D13A and L15A mutants exhibited secondary structure transitions into mixed population of both α- and β-structures in fibrils (Figures 2-3, summarized in Table 1). We speculate that the substitution of Lys9, Lys12 and Asp13 to alanine disrupted the formation of intra-helical salt-bridges between residues located 3-4 positions apart on the sequence, hence reducing α-helical stability (Kumar and Nussinov, 2002; Laabei et al., 2014). Phe11, Leu15, and especially Leu7, are central in forming the inter-sheet hydrophobic core observed in the cross-α structure (Figure 4), and their substitution to alanine likely to affect the propensity to form cross-α fibrils. This generally identifies the PSMα3 sequence as that which dictates the ability to switch: While the WT sequence stabilizes the cross-α fibrils, single point mutants shift the propensity to a mixture of α/β fibrils, and short truncations form strictly β-rich fibrils (Salinas et al., 2018). The chameleon properties embedded within the PSMα3 sequence are in common with other peptides in which helicity plays roles in fibril formation (Kim et al., 2016). It is nevertheless important to note that interactions with cell membranes can stabilize the helical configuration of PSMα3 (Malishev et al., 2018), and thus, the structural instability observed for the mutants in-vitro might be less significant upon peptide contact with cells.

The structural variations between PSMα3 mutants might affect the binding of the amyloid indicator dye ThT, which can explain the lack of ThT fluorescence for many of the mutants (L7A, F10A, F11A, K12A and D13A) (Figure S2), despite clear fibril-forming ability visualized by TEM (Figure 1A). Failure to bind ThT was previously reported for amyloid fibrils (Cloe et al., 2011; Nilsson, 2004), and was attributed to alteration in cavities along the fibril (Biancalana and Koide, 2010; Groenning et al., 2007). Hence, this assay is not sufficient to rule out fibril formation. The absence of ThT binding to PSMα3 mutants indeed led to the misinterpretation that these mutants do not form fibrils, and that toxicity is independent of fibrillation (Zheng et al., 2018). The results presented here further stress the importance of integration of methods in the evaluation of amyloid properties.

The presented observations in this manuscript suggest that membrane disruption and cytotoxicity of PSMα3 are regulated by fibrillation-dependent intra- and inter-helix electrostatic interactions. The complete lack of cytotoxicity of the K17A mutant, in contrast to other mutants in lysine residues that retained some cytotoxic activity (Figure 1B), corresponds with the absence of potential electrostatic interactions involving Lys17 within the fibril. In fact, K17A is the only alanine substitution for which we detected α-helical fibrils with no additional β-rich features, both in the FTIR spectra and fiber diffraction pattern (Figures 2-3), similar to the WT PSMα3, suggesting that Lys17 is not directly involved in stabilizing the cross-α fibrils. This putatively allows the free positive lysine charge within the cross-α fibril to interact with the cell membrane. Interactions between PSMα3 and membrane lipids lead to toxicity, as shown with the cross-α forming, all-D-amino-acid PSMα3 (D-PSMα3), which exhibited a similar level of cytotoxicity as WT PSMα3 (Figures 1&3), as was also recently reported by others (Yao et al., 2019). This lack of enantiomer stereo-specificity in PSMα3 toxicity advice against a mechanism mediated via direct interactions with membrane proteins, which are chiral. Aside from the lipids, additional membrane components, particularly in lipid rafts, such as cholesterol, glycoproteins and glycolipids, may also be involved in peptide-membrane interactions (Habchi et al., 2018; Hebda and Miranker, 2009; Laabei et al., 2014; Stefani, 2010; Walsh et al., 2014).

The activity of PSMα3 involves reciprocal interactions with cell membranes (Malishev et al., 2018). Specifically, eukaryotic membrane mimetics accelerate PSMα3 fibrillation, and the fibrils, in turn, affect the membranes and penetrate deep into the lipid bilayers (Malishev et al., 2018). This corresponds to other membrane-deforming peptides whose aggregation is often induced, or accelerated, by the membrane itself (Kinnunen, 2010; Pokorny et al., 2002; Zhang et al., 2014). Of note, the presence of lipids (interestingly those better mimicking bacterial cells) induced aggregation of the F3A mutant (Malishev et al., 2018). Thus, the environmental conditions, and particularly the presence of cells, might elicit fibril formation of otherwise non-fibrillating mutants, accounting for their reduced but not abolished toxicity. An earlier work on pathological amyloids suggested a process of fibril formation on membrane surfaces (Engel et al., 2008; Mahul-Mellier et al., 2015). For example, α-synuclein fibrils were shown to interact with the plasma membranes, and act as seeds, where the addition of monomers allows continuous aggregation within the membrane and increased toxicity (Mahul-Mellier et al., 2015). Co-aggregation of toxic amyloids with lipids was also supported by in-vivo studies of human α-synuclein (Gai et al., 2000; Sharon et al., 2003). This corresponds with our observation that aggregates of PSMα3 were co-localized with the membranes of affected cells (Figures 6&8). This putative dynamic co-aggregation process would require a mixed population of monomeric and fibrillar PSMα3 (Figure S6), whereas stabilized mature fibrils lacking equilibrium with soluble species would be less toxic. This might explain the reduced toxicity of an incubated racemic PSMα3 mixture, which showed robust fibrillation compared to the highly toxic L- and D-PSMα3 enantiomers (Yao et al., 2019). Nevertheless, the reduced toxicity of the racemic mixture (Yao et al., 2019) could also result from differences in the packing of the hydrophobic core and in the electrostatic interactions along the fibril.

Overall, our work elucidated key determinants governing PSMα3 cytotoxicity, namely positive charges, and cross-α fibrillation that regulates electrostatic interactions within the fibril and with the membrane, leading to membrane rupturing. Toxicity of pathological amyloids is often attributed to the prefibrillar and soluble oligomeric conformations, which often contain α-helices (e.g., (Ghosh et al., 2015)), including in initial stages of interactions with lipid membrane surfaces (Kinnunen, 2010), while the mature β-rich fibrils detoxify the amyloid (Stefani, 2012). Our study suggests that the process of cross-α fibrillation enhances toxicity, which may be related to the retained helical nature of its different species, as PSMα3 lacks secondary structure transition and remains helical upon aggregation (Figures 2-4). Based on the packing of the α-helices in the crystal structure of WT PSMα3 and three of its mutants (Figure 4), we expect that the initial aggregation of these amphipathic peptides involves interactions via their hydrophobic face to form helical bundles, which will eventually elongate to form the extensive hydrophobic core between the α-helical mated sheets. Accordingly, initial aggregates, or soluble oligomers, of PSMα3, should be similar in nature to the cross-α configuration. We overall hypothesize that toxicity is not necessarily attributed to a specific ‘toxic entity’, such as monomers, oligomers or fibrils, but is a ‘dynamic process’, in which a mixture of PSMα3 species and membrane lipids interact, affect each other, and co-aggregate, leading to membrane deformation.

## Acknowledgments

We acknowledge the guidance and technical support provided by Yael Pazy-Benhar and Dikla Hiya at the Technion Center for Structural Biology (TCSB), Yaron Kauffmann from the MIKA electron microscopy center of the department of material science & engineering at the Technion, Israel, the Russell Berrie Electron Microscopy Center of Soft Matter at at the Technion, Israel, Einat Nativ-Roth and Alexander Upcher from the Ilse Katz Institute for Nano-Science and Technology at Ben-Gurion University of the Negev, Israel, Einat Zelinger from The Interdepartmental Equipment Facility of the faculty of agriculture, food and environment, of the Hebrew university of Jerusalem, Israel, and of Nitsan Dahan and Yael Lupu-Haber from the LS&E infrastructure center of the Lokey center of the Technion, Israel. We are grateful for the service provided by Lihi Shaulov from the Rappaport Faculty of Medicine of the Technion, Israel, and by Na’ama Koifman from the Technion Center for Electron Microscopy of Soft Matter of the Russell Berrie Nanotechnology Institute, Israel. This research was supported by Israel Science Foundation (grant no. 560/16), Israel Ministry of Science, Technology & Space (grant no. 78567), DFG: Deutsch-IsraelischeProjektkooperation (DIP) (grant no. LA 3655/1-1), University of Michigan – Israel Collaborative Research Grant, BioStruct-X, funded by FP7, and the iNEXT consortium of Instruct-ERIC. The synchrotron MX data collection experiments were performed at beamlines ID23-EH2, ID29, and MASSIF-3 (ID30A-3) at the European Synchrotron Radiation Facility (ESRF), Grenoble, France, and at beamline P14, operated by EMBL Hamburg at the PETRA III storage ring (DESY, Hamburg, Germany). We are grateful to the teams at ESRF and EMBL Hamburg.

## Author Contributions

E.T-F. conceived, designed and conducted the experiments, determined the crystal structures and prepared figures; N.S. conducted experiments and prepared figures. O.T. designed several mutants and initiated toxicity experiments. E.T-F. and M.L wrote the paper with contributions from all authors.

## Supporting Figures and Table

**Figure S1.**
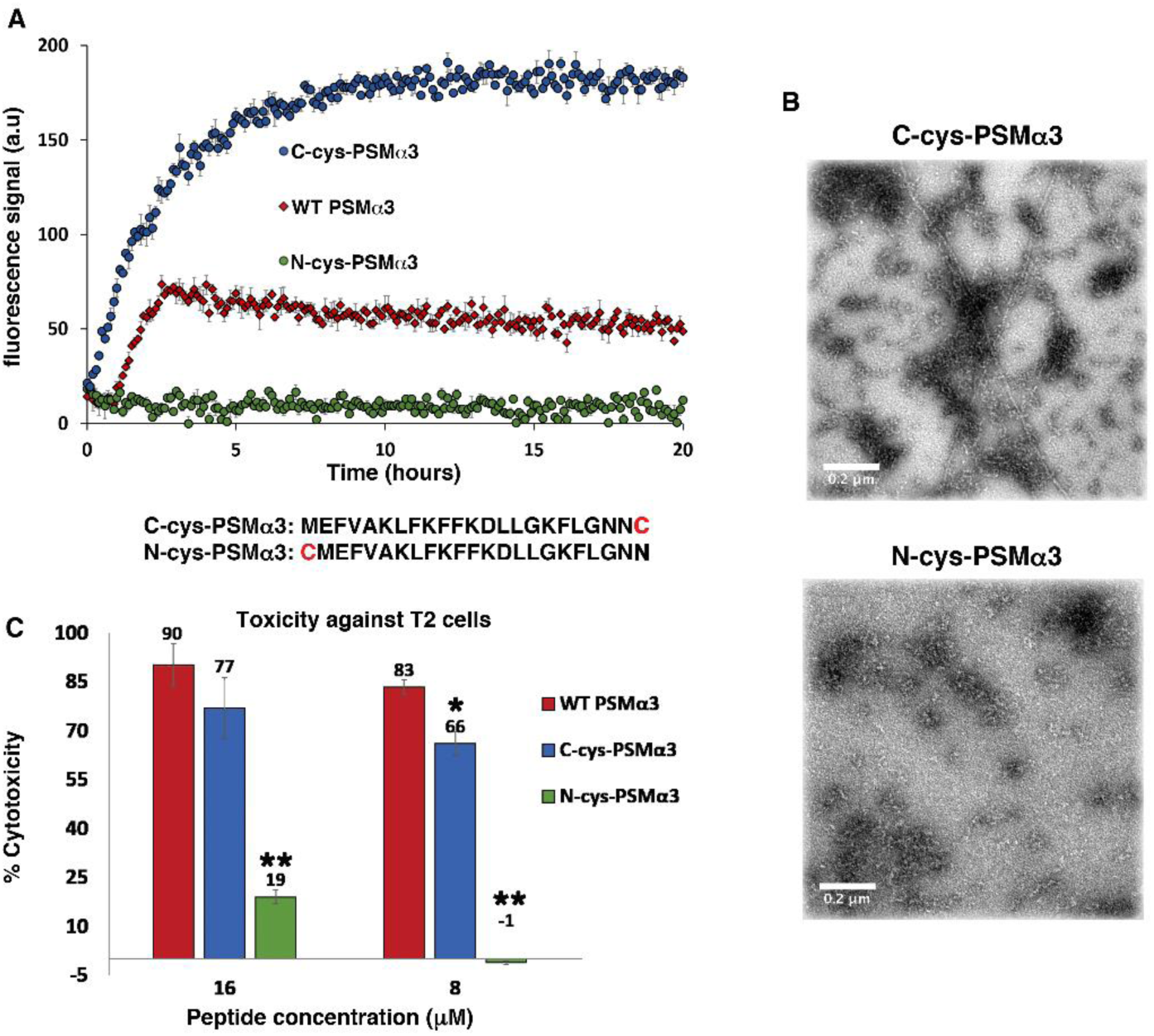
Fibrillation and cytotoxicity of cysteine derivatives of PSMα3. Fibrillation and cytotoxicity of PSMα3 derivatives with cysteine residues added to either the N-terminus (N-cys-PSMα3) or C-terminus (C-cys-PSMα3); sequences are depicted. (**A**) ThT fibrillation kinetic curve showing the fast fibrillation rate of C-cys-PSMα3 compared to WT PSMα3, and almost no ThT binding to N-cys-PSMα3. (**B**) TEM micrographs showing fibrils of C-cys-PSMα3, while no fibrils were found for N-cys-PSMα3. (**C**) C-cys-PSMα3 is toxic to T cells while N-cys-PSMα3 showed significantly reduced cytotoxicity. The experiment was performed in triplicates and repeated three times, on different days. A representative graph is presented. Error bars represent standard error of the mean of each triplicate. * p<0.05 and ** p<0.001 were calculated using the Mann-Whitney U (two tailed) test in comparison to the values of WT PSMα3, accounting for the data from all three repeats.

**Figure S2.**
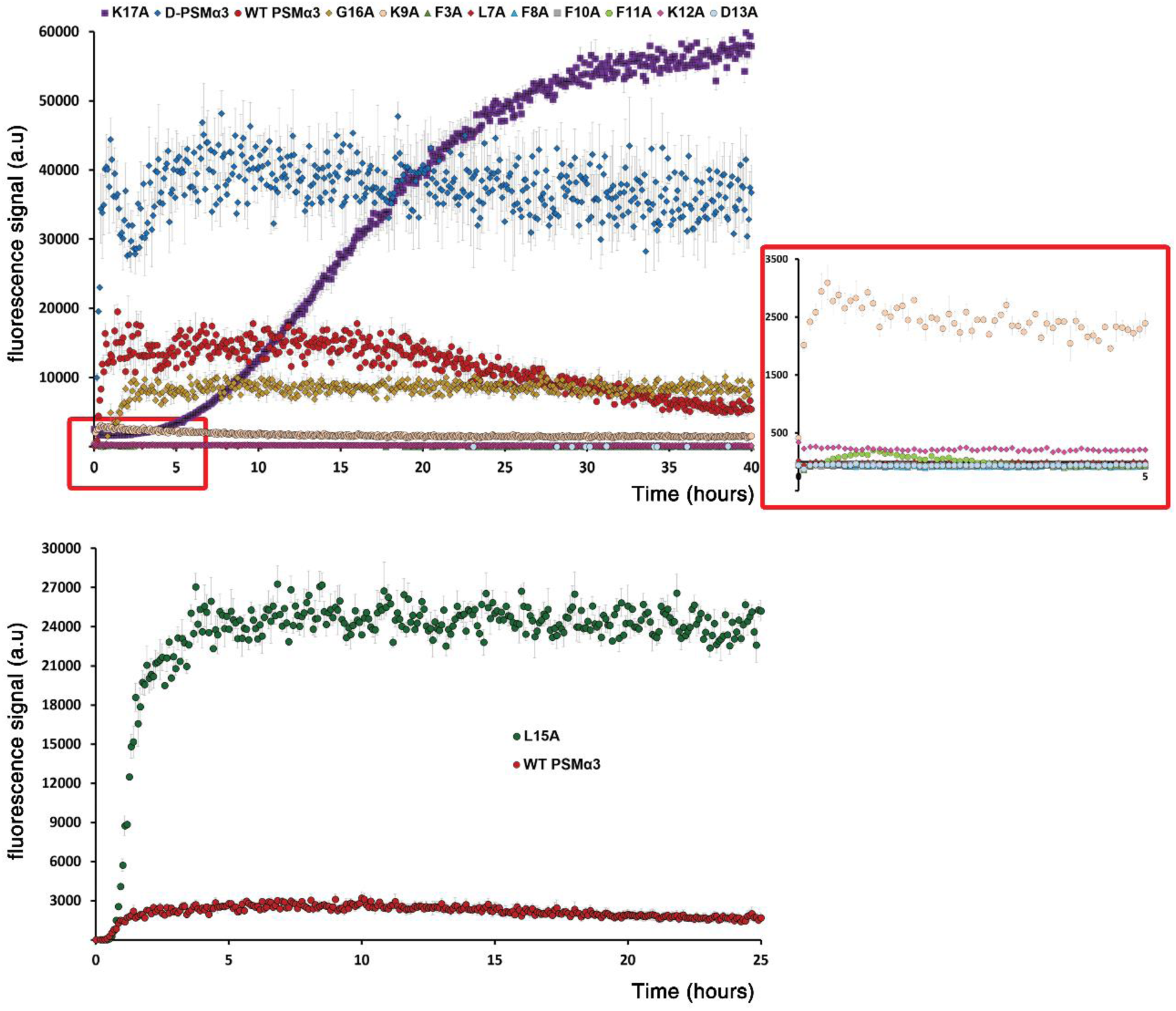
Fibrillation kinetics of PSMα3 mutants. ThT binding to PSMα3 derivatives shows the typical fibrillation kinetics, which includes a lag time followed by aggregation. The peptides (50 µM) were mixed with ThT (200 µM) in a buffered solution (pH 8). WT PSMα3, as well as D-PSMα3, G16A, and K17A mutants bound ThT with different kinetics (top left). The inset on the right presents a weaker binding of ThT to the K9A mutant, but nevertheless clear fibrillation kinetics were measured. L15A was associated with an especially high ThT signal and its fibrillation curve is therefore depicted separately from the others (bottom). Overall, PSMα3 and five derivatives were shown to bind ThT, while L7A, F10A, F11A, K12A, and D13A did not bind ThT, but did form fibrils visualized by TEM (Figure 1).

**Figure S3.**
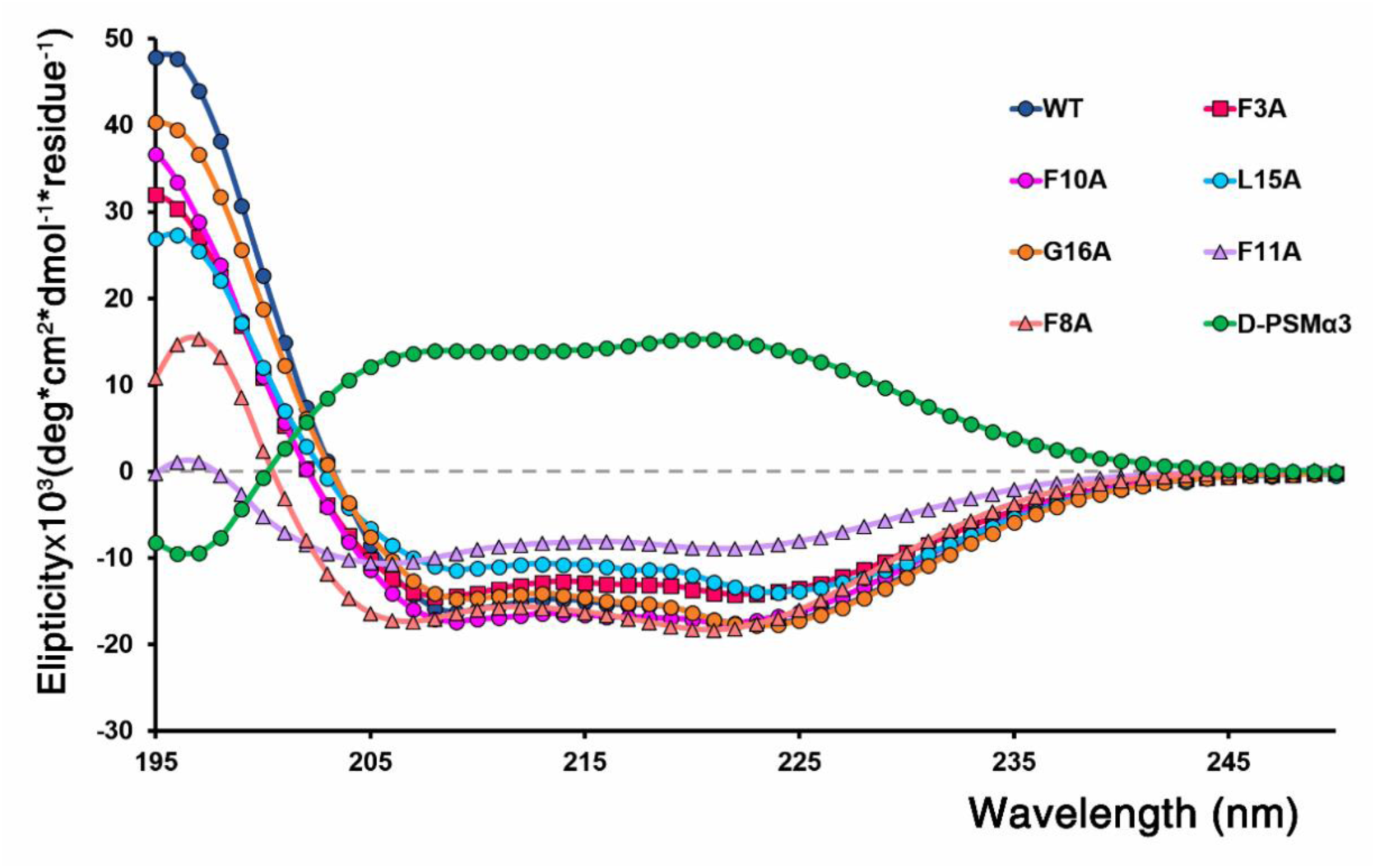
Secondary structure analysis of WT PSMα3 mutants in solution. The circular dichroism (CD) spectra of PSMα3 mutants showed that they maintained α-helicity, as depicted by the double minima near 208 nm and 222 nm typical of a α-helical conformation. F8A and F11A exhibited somewhat reduced helicity but were still mainly helical in solution. The measurements were done using 111 µM peptides in a buffer containing 150 mM sodium chloride and 10 mM sodium phosphate at pH 7.4. The concentrations of the peptides were calculated during the measurement to account for the effect of aggregation, as described in the Experimental section.

**Figure S4.**
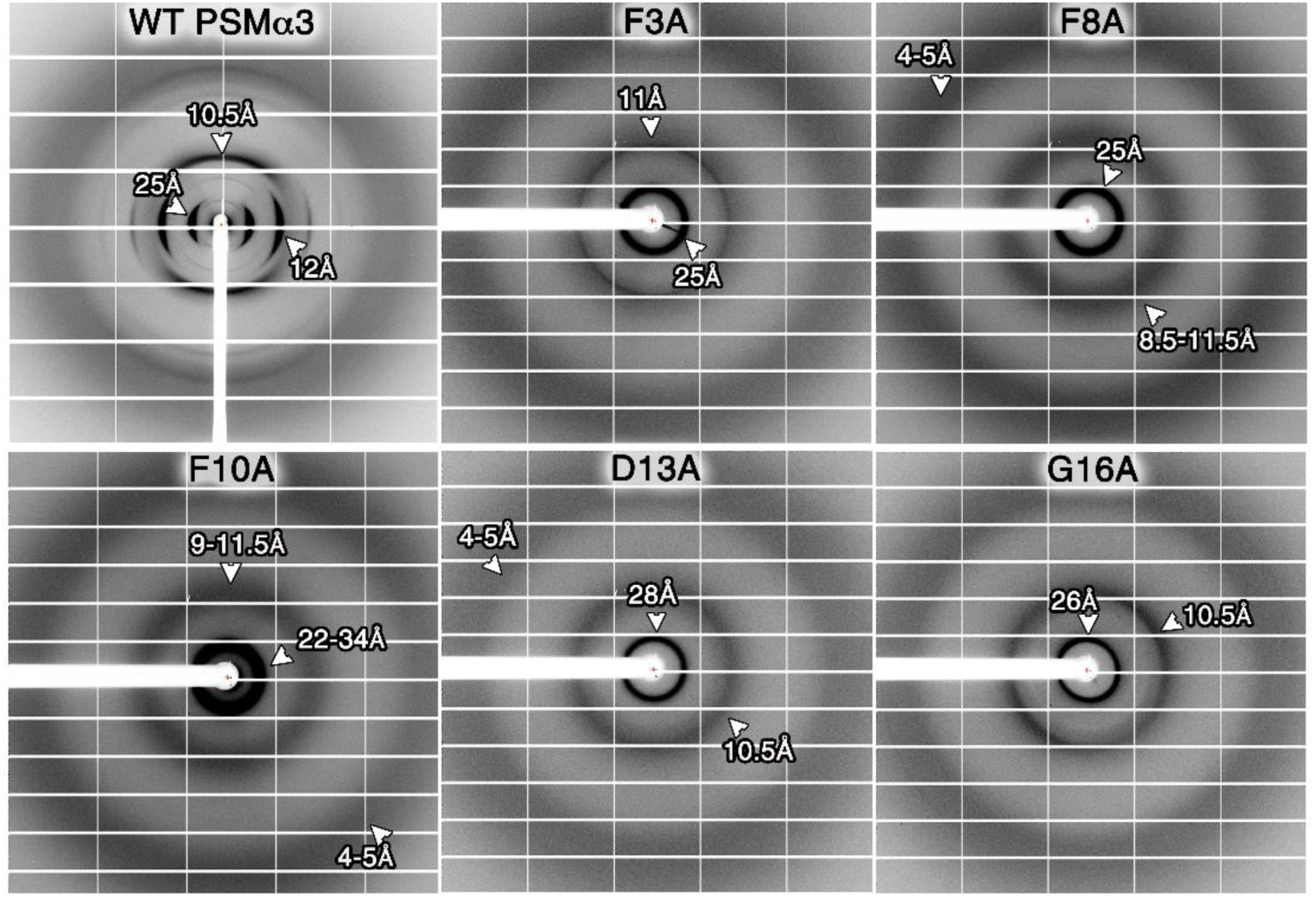
X-ray fiber diffraction of PSMα3 mutants which showed diffuse patterns. X-ray fiber diffraction of WT PSMα3 exhibited a cross-α pattern. The fiber diffraction of the mutants F3A, F8A, D13A, and G16A all showed diffuse fiber diffraction, hindering definitive interpretation of the data, despite repeated attempts. Main reflections, although diffuse, are depicted. Of note, in contrast to WT PSMα3 that formed clear fibrous species on the edge of the capillary when prepared for fibril diffraction, the non-fibrillating mutants, F3A and F8A, did not form any clear fibrous structures; thus, X-ray data were collected from a dried sample that remained on the capillary.

**Figure S5.**
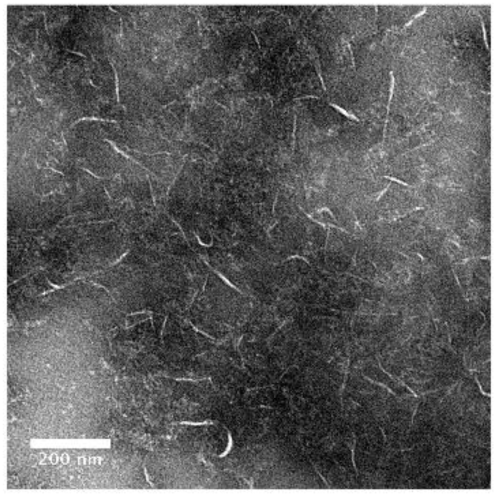
FITC-labeled PSMα3 forms fibrils. The micrograph shows fibrils formed by FITC-labeled PSMα3. FITC-PSMα3 was dissolved in ultra-pure water to 10 mM and incubated at RT for 7 days. Fibrils were then separated from the supernatant, diluted to 1mM and deposited on a carbon-coated grid. A 200 nm scale bar is depicted.

**Figure S6.**
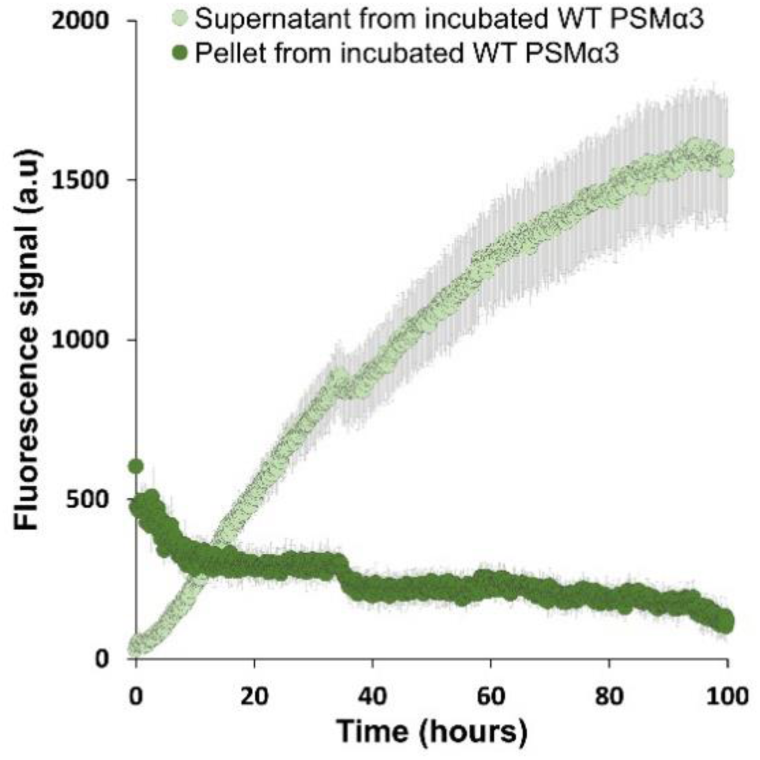
Dynamics in WT PSMα3 fibril assembly/disassembly. Supernatant separated from an incubated (3 days at 37ᵒC) 10mM WT PSMα3 solution (with theoretically calculated concentration of 50µM after dilution) contained peptide species that could further aggregate and bind ThT (200µM) to demonstrate a canonical aggregation kinetics curve. The pellet from this incubated PSMα3 sample (also 20-fold diluted) exhibited a relative plateau (some decrease in signal might be due to fibril dissociation). The experiment was performed with triplicates in two repeats, a representative curve is presented. Error bars represent a standard error of the mean from a triplicate.

**Table S1.** Calculated distances in the structures of WT PSMα3 and mutants. The dry interface corresponds to the hydrophobic packing between mated sheets. The amphipathic nature of the peptides results in alternating rows of dry and wet interfaces in the crystal packing. In the wet interface, interactions between sheets are mediated by water molecules. The distances were calculated in Chimera (Pettersen et al., 2004) using a different method than in our previous report (Tayeb-Fligelman et al., 2017), therefore resulting in a slightly different inter-sheet value for the WT PSMα3.

| Peptide | | WT-PSMα3 | L15A | G16A | K12A |
| --- | --- | --- | --- | --- | --- |
| Inter-sheet distances (Å) | Dry interface | 12.6 | 13.0 | 10.2 | 10.2 |
|  | Wet interface | 15.6 | 13.6 | 13.9 | 13.9 |
| Inter-helix distance along each sheet (Å) |  | 10.5 | 10.6 | 13.1 | 13.0 |
| Length of α-helical axis | | 25.5 | 26.9 | 25.1 | 24.9 |

## Experimental Section

Further information and requests for resources and reagents should be directed to and will be fulfilled by Lead Contact Meytal Landau.

**Table 3.**
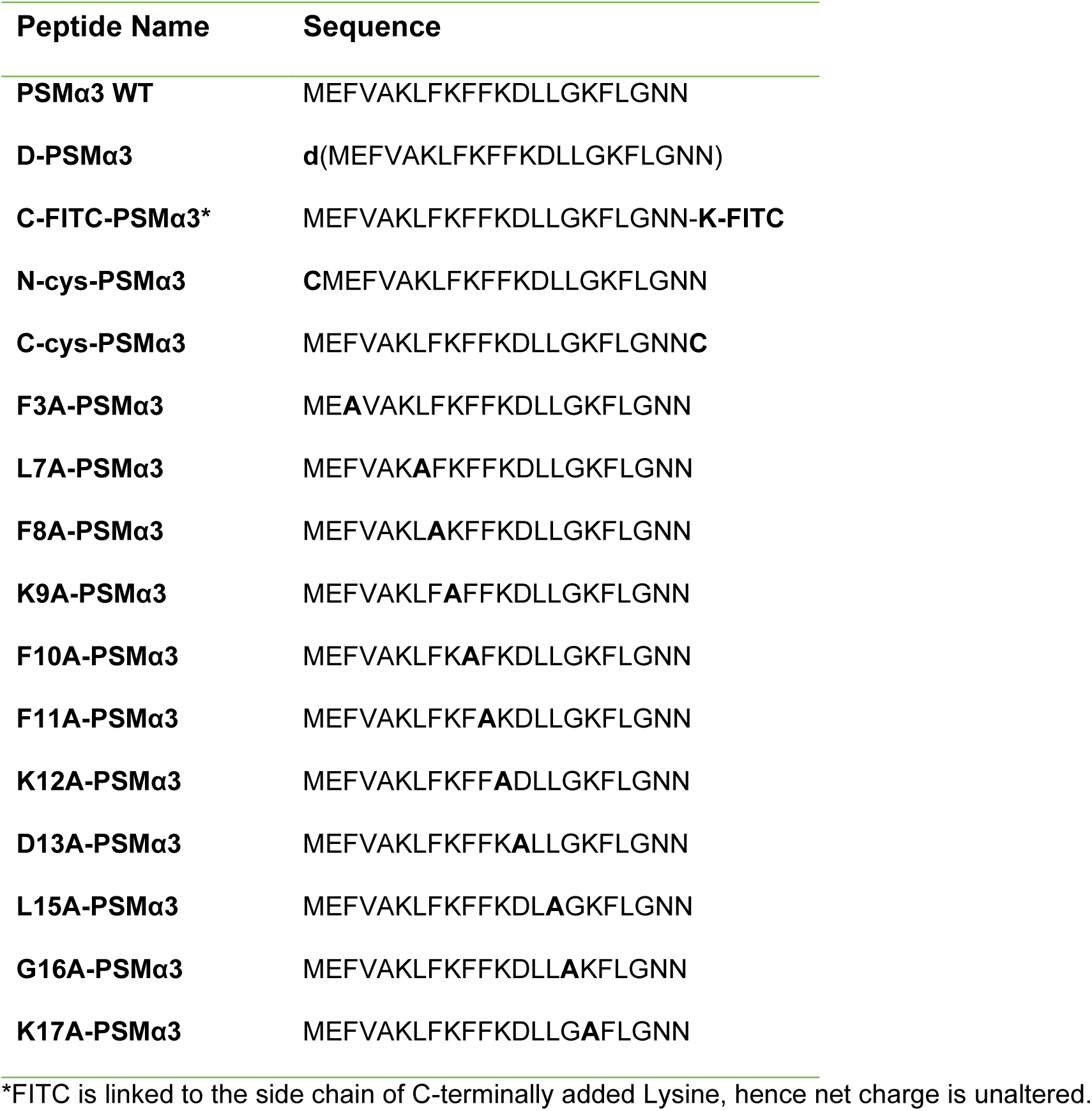
List of PSMα3 derivatives.

PSMα3 WT and mutant peptides (custom synthesis at >98% purity) were purchased from GL Biochem (Shanghai, China) Ltd. or from GenScript. C-FITC-labeled PSMα3 was purchased from GL Biochem. All chemicals and reagents were of analytical grade. Chemicals were purchased from Sigma-Aldrich Ltd., including: trifluoroacetic acid (TFA), hexafluoroisopropanol (HFIP), thioflavin T (ThT), and deuterium-oxide (D2O). Ultra-pure water and Dullbeco’s phosphate buffered saline (PBS) were purchased from Biological Industries (Kibbutz Beit-Haemek, Israel). Phosphate dibasic anhydrous and di-sodium hydrogen phosphate were purchased from Carlo Erba, distributed by Romical Ltd. (Beer-Sheva, Israel). Sodium chloride was purchased from Merck. Additional reagents and consumable are mentioned below.

### Peptide pretreatment for disaggregation

In experiments that included a time-resolved aspect or secondary structure determination in different peptide states, for example measuring aggregation kinetics, and CD and FTIR measurements, the peptides were pretreated to disassemble any pre-formed aggregates, and to ensure reproducibility of the results (Chen and Wetzel, 2001; Jao et al., 1997; O’Nuallain et al., 2006). Lyophilized peptide powder was freshly dissolved in TFA-HFIP (1:1), to a concentration of 1-5 mg/ml, sonicated for 5-10 min in a sonication bath, and then allowed to evaporate under vacuum or in a chemical hood, for 2-3 days. Residual solvent was evaporated using a speed vac. For certain experiments, the peptides were incubated for 1h in TFA-HFIP at room temperature (RT) and dried directly in an acid-resistant speed vac (RVC 2-18 CDplus, CHRIST, distributed by Dr Golik, Tel-Aviv, Israel). Unless immediately tested, the treated peptides were stored at −20°C.

### Fibrillation assays

#### Thioflavin T fluorescence kinetic assay

Fresh ThT reaction solution was prepared by diluting (1:10) the filtered 2 mM ThT stock, made in ultra-pure water, into the reaction buffer containing 10 mM sodium phosphate buffer and 150 mM NaCl, pH 8.0. PSMα3 and its derivatives (Table 3), pretreated as described above, were dissolved in ultra-pure water to 1 mM, on ice, and sonicated for 5 min using a sonication bath. The peptides were then diluted, on ice, in a filtered reaction buffer, and centrifuged (5 min, 4 °C, 10,000 rpm). Centrifugation of K17A was avoided as only the non-centrifuged K17A samples showed ThT binding. The supernatant of each mutant (or a sample of K17A solution) was separately mixed with ThT reaction solution in the wells of a Greiner bio-one black, 96-well, flat-bottom plate (distributed by De Groot Medical Ltd. Rosh Ha’ain, Israel), to final concentrations of 50 µM peptide and 200 µM ThT. The plate was immediately covered with a silicone sealing film (ThermalSeal RTS, Sigma-Aldrich), and incubated in a plate reader (CLARIOstar or FLUOstar Omega, BMG LABTECH. Distributed by Zotal Ltd. Tel-Aviv, Israel) at 37 °C, with 500 rpm shaking for 15 sec before each cycle, up to 1000 cycles of 3-5 min each. ThT fluorescence was measured at an excitation wavelength of 438±20 nm and emission wavelength of 490±20 nm. Measurements were made in triplicates and each experiment was repeated at least three times. All triplicate values were averaged, appropriate blanks were subtracted, and the resulting values were plotted against time. Calculated standard errors of the mean from a triplicate are presented as error bars. Of note, the measurement of L15A was separated from the measurement of the other mutants since L15A produced a very high signal which caused signal overflow and required digital gain reduction in the plate reader. Cysteine derivatives (originally designed for downstream fluorescent labeling) were also tested in a separate experiment (Figure S1), including PSMα3 for comparison, and according to the exact same protocol.

The assembly/disassembly properties of fibrillated PSMα3 (Figure S6) was assessed as follows: lyophilized (untreated) PSMα3 powder was dissolved in ultra-pure water to a concentration of 10mM and incubated for 3 days at 37°C to allow fibril formation. The incubated sample was centrifuged at 14,800xg for 20 minutes and the supernatant was separated from the pellet, which was re-suspended to a similar final volume in ultra-pure water. Both separated supernatant and pellet samples were diluted 200-fold with the above reaction buffer containing 200 µM of ThT. The reaction was carried out according to the above protocol with the exact same parameters.

#### Transmission electron microscopy (TEM)

Lyophilized (untreated) peptide powders were separately dissolved in ultra-pure water to a concentration of 10 mM, and incubated for 7-14 days at RT, or for 3-7 days at 37 °C. All samples were then centrifuged for 20 min at 14,800 rpm. The supernatant was discarded, and the pellet was resuspended to the same volume and diluted 10-fold in ultra-pure water. Samples (5 µl) were then directly applied onto 400-mesh copper TEM grids with support films of Formvar/Carbon (Ted Pella, distributed by Getter Group Bio Med, Petah-Tikva, Israel), that were charged by high-voltage, alternating current glow-discharge (PELCO easiGlow, Ted Pella), immediately before use. Grids were allowed to adhere for 2 min. Excess solution was blotted with filter paper, and the grids were negatively stained with 2% uranyl acetate for 30 sec, blotted again, and dried under vacuum. Specimens were examined with a FEI Tecnai T12 G2 or FEI Tecnai G2 T20 S-Twin transmission electron microscope (FEI, Hillsboro, Oregon, United States), at an accelerating voltage of 120 or 200 KeV, respectively.

### Circular dichroism

All peptides were pretreated with TFA: HFIP 1:1 mixture as described above in a concentration of 1 mg/ml in, in order to disassemble any pre-formed aggregates, and then fully dried. The treated peptides were dissolved in ultra-pure water to 1.5 mM and then sonicated for 10 min, in a sonication bath. For each derivative tested, 250 µl of 10 mM sodium phosphate, 150 mM NaCl pH 7.4 buffer was first measured as a blank, after which, 20 µl peptide was diluted into the cuvette (to a calculated concentration of 111 µM). Samples were thoroughly mixed to ensure homogeneity immediately prior to measurement. CD spectra were recorded on either the Applied Photophysics Chirascan qCD or Pi-star 180 (Applied Photophysics, Surrey, United Kingdom), using a 1 mm path-length fused quartz cell. The presented measurements are an average of four scans for each peptide or two scans for blank, captured at a scan speed of 1 sec, bandwidth and step size of 1 nm, over a wavelength range of 180-260 nm. High-voltage (HV) and absorbance measurements were taken simultaneously, and CD values with corresponding HV or absorbance values above the recommended limit (evident only in wavelengths shorter than 200 nm), were omitted from the spectra. Due to peptide aggregation upon dilution in the tested buffer, which significantly lowered the soluble peptide concentration for certain PSMα3 derivatives, the concentration of each sample at real time was experimentally determined by UV absorbance, which was measured in parallel to CD spectra measurements, in the same machine. Since PSMα3 sequence lacks tryptophan and tyrosine residues, the soluble peptide concentration (in molar units) was calculated using the Beer-Lambert equation at 205 nm: *A*_205_ = ε _205_ × *l* × *C*. The pathlength l = 0.1 cm. A205 was determined by averaging the blank corrected absorbance values at 205 nm from all four scans. ε205 (M^-1^ x cm^-1^) is a sequence-specific extinction coefficient calculated using the webserver (http://spin.niddk.nih.gov/clore/) (Anthis and Clore, 2013). The molar ellipticity per residue (Ɵ, in mdeg*cm^2^ *dmol^-1^*residu^-1^) was derived from the raw data (δ, in mdeg.), using the following formula: Ɵ = δ / (L*C*N), where L is the path length of the cuvette, C is the real-time calculated molar concentration of PSMα3, and N is the number of residues in PSMα3 (i.e., 22).

### ATR-FTIR spectroscopy

Attenuated total-internal reflection Fourier transform infrared (ATR-FTIR) spectroscopy was conducted to test secondary structure features in the fibril form. PSMα3 and derivatives, pretreated as described above, were separately dissolved in 5 mM HCl to 1 mg/ml, sonicated for 3 min in a sonication bath, frozen in liquid nitrogen and lyophilized. Upon complete dryness, the peptide was subjected to the same treatment but without sonication. The latter step was repeated four times in order to replace all TFA molecules that served as counter ions in the synthesis of the peptide and that were added during the pretreatment. The dry peptide was then dissolved in D2O to 1 mg/ml, frozen and lyophilized. This process was repeated at least twice. The dry peptide was further dissolved in D2O to 25 mg/ml and incubated at RT for 2-6 h. Sample (5 µl) were then separately applied on a surface of the ATR module (MIRacle Diamond w/ZnSe lens 3-Reflection HATR Plate; Pike Technologies, Fitchburg, Wisconsin, USA) and left to dry under nitrogen, to form a thin film. Absorption spectra were recorded with a Tensor 27 FTIR spectrometer, equipped with a high-sensitivity LN-cooled MCT detector (Bruker Optics, Bruker Scientific Israel Ltd., Rehovot, Israel). Measurements were performed as an accumulation of 100 scans with 2 cm^-1^ resolution over a range of 1500-1800 cm^-1^. The same procedure was repeated after incubation of samples for seven days, at RT, in a sealed tube. The residual samples were then centrifuged for 10 min at 17,000g, at RT, to separate the fibrils from the supernatant. The pellet was resuspended with the same amount of D2O, centrifuged again under the same conditions, and the remaining pellet was resuspended in 10 µl D2O and measured under the same conditions. Background (ATR module) and blank (D2O) subtraction, and baseline correction, was performed using the Bruker OPUS software. ATR-FTIR curves were normalized to unity and their second derivatives were calculated and rendered using the Origin 2018b graphing & analysis software (OriginLab^®^, distributed by Tashtit Scientific Consultants Ltd, Rishon LeZion, Israel). The amide I’ region of the spectra (1600-1700 cm^-1^) is presented in the curves.

### Tissue culture and cell toxicity

Our calibration experiments verified that T2 cytotoxicity results were not affected by peptide pretreatment, therefore, in this work, cell toxicity assays were conducted with the non-treated peptide powders. T2 cells (174 x CEM.T2) (ATCC^®^ CRL-1992™, distributed by Biological Industries) were cultured in RPMI 1640 medium containing L-glutamine (Gibco, distributed by Rhenium Ltd. Modi’in, Israel), supplemented with 20% heat-inactivated fetal calf serum, penicillin (100 U/ml) and streptomycin (0.1 mg/ml), at 37°C in 5% CO2. Before each experiment, the cells were washed and resuspended in assay medium, which was similar to T2 growth medium, except that it contained 0.5% fetal calf serum. The cells were incubated for two hours at 37 °C, in 5% CO2. PSMα3 and its derivatives were separately dissolved to 1 mM in ultra-pure water, sonicated for 10 min, divided to aliquots and stored at −20°C. One aliquot was thawed immediately before each experiment. Peptides were serially diluted in assay medium to double the final concentration used in the assay (as denoted in the figures). Then, 50 µl of each diluted sample was pipetted into a well of a 96-well plate in triplicates (or duplicate for one repeat of the experiment for technical reasons), and incubated at 37 °C, in 5% CO2, without shaking, for 1 h. Two 96 well plates were used per experiment to accommodate all mutants while testing different peptide concentrations. WT PSMα3 was present in both plates in each repeat of the experiment, hence doubly measured in comparison to the mutants, and the % cytotoxicity reported is the average from both plates for each experiment. Cells were then diluted in assay medium to 0.15×10^6^ viable cells/ml and 50 µl were transferred to each well using a multichannel pipette. The plate was incubated for 30 min, at 37 °C, in 5% CO2. Cell death and cell lysis were quantified using the LDH colorimetric assay, according to the manufacturer’s instructions, including all recommended controls (LDH; Cytotoxicity Detection Kit Plus, Roche Applied Sciences, Penzberg, Germany). Cell-free culture medium was measured as background. Cells subjected to the same experimental conditions apart from peptide addition, were used as the ‘low control’ to account for spontaneous LDH release. Cells subjected to the same experimental conditions apart from peptide addition, and treated with lysis buffer prior to color development reaction, according to the manufacturer instructions, were used as the high positive control to account for maximum LDH release. The average absorbance values of each replicate and controls, measured at 490 nm with a reference wavelength at 690 nm using the FLUOstar Omega (BMG LABTECH) plate reader, was calculated, and background was subtracted. The following equation was used to calculate % cytotoxicity according to the kit’s protocol: 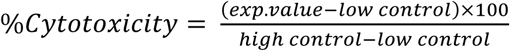. The data were analyzed using Excel. The experiments were repeated three times on different days. The PSMα3 cysteine derivatives were also examined in separate experiments and compared with WT PSMα3. Statistical analysis was conducted in XLSTAT 2018 software (Addinsoft) using the Mann-Whitney U (two tailed) test. Sample sizes for the toxicity assay were n=16 for WT PSMα3, and n=8 for all mutants apart from F11A for which n=7 due to one outlier. The sample size was n=9 for all peptides tested in the toxicity assay of the cysteine derivatives (fig. S1), accounting for a triplicate from all three repeats.

### Confocal microscopy

All solutions used for the confocal microscope assays, apart for cell media, were prepared with filtered DDW and/or were filtered twice using a 0.22 µm syringe filter.

#### PSMα3 aggregation on the membranes of fixed cells

Untreated WT and FITC-labeled PSMα3 peptides were separately dissolved in ultra-pure water to a concentration of 2 mM, mixed to a final concentration of 1 mM, and then incubated together at RT just prior to the experiment. T2 cells (174 x CEM.T2) (ATCC^®^ CRL-1992™) grown as described above, were washed and diluted in unsupplemented RPMI 1640 medium (Gibco) to 5×10^6^ viable cells/ml and gently pipetted onto dry, small-diameter coverslips pre-coated with 0.1 % poly-lysine solution, in a 24-well plate. Samples were incubated on the coverslip for 20 min, at 37 °C, in 5% CO2. In parallel, the peptide mixture was diluted in the cytotoxicity assay medium, comprised of RPMI 1640 medium supplemented with 0.5 % heat-inactivated fetal calf serum, penicillin (100 U/ml) and streptomycin (0.1 mg/ml), to a final concentration of 5 µM of each (WT and FITC-labeled peptides). The cell-coated coverslips were then washed three times with fresh, pre-warmed (to 37 °C) and unsupplemented RPMI medium, and either cytotoxicity assay medium alone (for control), or the above described peptide mixture were applied on the coverslips. The cells were incubated for 10 min at 37 °C in 5% CO2, rinsed three times with filtered PBS (Biological Industries), and fixed at RT, for 1 h, in aqueous solution containing 100 mM sucrose, 100 mM sodium phosphate buffer, pH 7.4, 2.5% glutaraldehyde (from a 8% stock solution, Electron Microscopy Sciences, Hatfield, Pennsylvania, USA), and 2% paraformaldehyde (from 16% stock solution, Electron Microscopy Sciences). After fixation, the cells were washed three times with filtered PBS. Of note, all solution exchange steps were carried out rapidly, to avoid any contact of the cells with air. Finally, the cells were immersed in freshly filtered PBS and stored at 4ᵒC until visualized with a Leica SP8 confocal laser scanning microscope, equipped with a HCPL APO CS 2 40x/1.10 water objective. The images were taken at a zoom of either 0.75 or 2.3. Treated cells were images at a Z stack series of either 20 slices of ∼0.5 µm each (for the 10min treated cells) or 109 slices of ∼0.08 µm each (for 30min treated cells). Scale bars were added and the Images were processed and rendered with ImageJ. A 3D image of the treated cells was reconstructed from each Z stack series using the “Z project” function in ImageJ to account for the entire cell and for a better comparison with the controls. The background signal was subtracted with the “rolling ball function” using the “sliding paraboloid” and “disable smoothing” options. Brightness and contrast levels of each constructed image were adjusted so to best visualize the features in the image, and finally the image was slightly cropped. It is of note that the experiment with fixated cells was performed only once, however the effect of the peptide on the cells was tested numerous times using live imaging microscopy, with different peptide-to-cell ratios and imaging periods, all showing aggregation of the peptide on the cell membrane in the initial interaction steps, prior to complete cell penetration and cell death (visualized with propidium iodide).

#### PSMα3 co-localization with cell membranes

On the day of the experiment, untreated WT and FITC-PSMα3 were separately dissolved in ultrapure water to a concentration of 1 mM and mixed together in PBS to concentrations of 10 µM and 20 µM, respectively. Wheat germ agglutinin (WGA) Alexa Fluor™ 350 Conjugate stock solution (ThermoFisher Scientific) was prepared in PBS to a concentration of 1 mg/ml and then diluted in Hank’s balanced salt solution (HBSS) to a concentration of 5 µg/ml, according to the manufacturer’s instructions. T2 cells (174 x CEM.T2) (ATCC^®^ CRL-1992™) were grown as described above in RPMI 1640 medium containing L-glutamine (Gibco), supplemented with 20% heat inactivated fetal calf serum, penicillin (100 U/ml) and streptomycin (0.1 mg/ml), at 37 °C in 5% CO2. Before the experiment, 1×10^6^ cells were washed and resuspended in assay medium, similar to T2 growth medium, except that it contained 0.5% fetal calf serum. After two hours of incubation at 37 °C, in 5% CO2, the cells were washed and suspended in the WGA staining solution and incubated for 10 min, under the same conditions, for membrane staining. The cells were then washed twice and diluted in assay medium to a density of 0.15*10^6^ cell/ml. Stained cells were transferred to a chamber of a 6-channel microscope µ-slide (µ-Slide VI 0.4, #cat 80606, ibidi) and imaged. The incubated peptide mixture was supplemented with propidium iodide from a stock of 1 mg/ml in PBS, and then injected to the µ-slide channel and allowed to diffuse toward the cells. Final concentrations in the chamber were 5 µM WT, 10 µM FITC-PSMα3, and 0.02 mg/ml PI. Cells were continuously imaged with an LSM 710, AnxioObserver laser scanning confocal microscope (SEM; Zeiss), equipped with a C-Apochromat 40x/120 W Korr M27 water objective, for 50 min, at 37 °C and 5% CO2. Data analysis was performed using the ZEN 2.3 lite (Zeiss) and a representative image was rendered with ImageJ. The thresholds of the different color channels were individually adjusted.

### Scanning electron microscopy

All solutions used for this assay, apart from cell medium, were prepared with filtered DDW and/or were filtered twice using a 0.22 µm syringe filter. SEM chips (silicon chips, 5×5 mm) were pre-coated with 0.1% poly-lysine solution, washed three times with DDW and dried. Untreated PSMα3 peptide was freshly dissolved in ultra-pure water to a concentration of 1 mM and sonicated for 5 min in a sonication bath, and then further diluted to a concentration of 10 µM, in cytotoxicity assay medium, comprised of RPMI 1640 medium supplemented with 0.5 % heat inactivated fetal calf serum, penicillin (100 U/ml) and streptomycin (0.1 mg/ml). As control, ultra-pure water was added to the medium in the same volume as the peptide. T2 cells (174 x CEM.T2) (ATCC^®^ CRL-1992™) grown in RPMI 1640 medium containing L-glutamine (Gibco), supplemented with 20% heat inactivated fetal calf serum, penicillin (100 U/ml) and streptomycin (0.1 mg/ml), were washed and diluted in unsupplemented RPMI medium to a concentration of ∼10×10^6^ viable cells/ml. The cell suspension was applied onto the poly-lysine coated chips, and incubated for 20 min. The chips were then rinsed three times with fresh RPMI medium, and immediately immersed, for 5 min, in either 10 µM PSMα3 or the control solutions. The chips were then rinsed twice in filtered PBS (Biological Industries). Then, the chips were incubated, for 1 h, at RT, in aqueous fixation solution containing 100 mM sucrose, 100 mM sodium phosphate buffer, pH 7.4, 2.5% glutaraldehyde (from an 8% stock solution, Electron Microscopy Science) and 2% paraformaldehyde (from 16% stock solution, Electron Microscopy Science). Chips were then washed in three 2-min cycles, with 0.1 M cacodylate buffer, pH 7.4, and incubated, for 15 min, at RT, in freshly prepared secondary fixative containing 1% OsO4 in 0.1 M cacodylate buffer. The chips were then subjected to three 2-min washing cycles with DDW, followed by dehydration by washing the chips twice in a row in each concentration of a graded ethanol series of 15%, 30%, 50%, 70%, 90%, 95% and 100% with 3 min incubation for each wash, apart from the last step which lasted 5 min. Finally, the chips were dried using the critical point drying (CPD) method, coated with 5 nm of chromium and kept under vacuum until they were scanned with a Zeiss ULTRA *plus* field emission SEM using either the InLens or SE2 detectors (Carl Zeiss, Oberkochen, Germany). This experiment was repeated once.

### Fiber X-ray diffraction

Peptides were dissolved in ultra-pure water, to a concentration of 20 mg/ml. Several microliters were applied between two sealed glass capillaries and incubated for up to 10 h, at RT. For some peptides, the X-ray fiber diffraction was collected immediately upon complete drying of the fibrils on the capillaries for obtaining clear diffraction pattern. For WT PSMα3, the diffraction pattern was tested using a freshly dissolved peptide, and with a peptide solution incubated in the tube for 10 h at RT, both showing a similar diffraction pattern. The peptides presented in Figure S4 yielded only diffuse diffraction patterns at all conditions tested. X-ray diffraction was collected at the micro-focused beam P14 at the high brilliance 3rd Generation Synchrotron Radiation Source at DESY: PETRA III, Hamburg, Germany or on beamlines ID23-EH2 and ID29 at the European Synchrotron Radiation Facility (ESRF), Grenoble, France.

### Peptide crystallization

Peptides were crystallized with no pretreatment. After dissolving the peptides, the solutions were sonicated in a sonication bath for 5-10 min, following by 5 min centrifugation at 13,700 rpm, at 4 ᵒC. The peptides were kept on ice until crystallization plates were set. All crystals were grown at 20 °C, using the hanging-drop vapor diffusion technique. K12A crystals were grown from a 10 mM (26.2 mg/ml) solution in ultra-pure water, mixed with a reservoir (Qiagen, Classics Suite I crystallization screen, condition B8) containing 0.1 M HEPES pH 7.5, 0.5 M ammonium sulfate, and 30% v/v 2-methyl-2,4-pentanediol (MPD). Crystals were flash-frozen before X-ray data collection. No cryo-protectant was added. L15A crystals were grown from a 7.5 mM (19.2 mg/ml) solution in ultra-pure water, in a reservoir (Molecular Dimensions, JCSG + HTS crystallization screen, condition E5) containing 0.1 M CAPS, pH 10.5 and 40% v/v MPD. Crystals were flash-frozen before X-ray data collection. No cryo-protectant was added. G16A crystals were grown from a 10 mM (26.2 mg/ml) solution in ultra-pure water, in a reservoir (Qiagen, Classics Suite I crystallization screen, condition A1) containing 0.1 M sodium acetate, pH 4.6, 0.01 M cobalt(II) chloride, and 1 M 1,6-hexanediol. Before data collection, the crystals were immersed in a cryo-protectant solution in which the ingredients of the crystallization reservoir were supplemented with 20% ethylene glycol, and then flash-frozen in liquid nitrogen.

### Structure determination and refinement

X-ray diffraction data of PSMα3 mutants were collected at 100 °K at the micro-focal beamlines ID23-2 (G16A) and ID30a-3 (K12A, L15A) at the European Synchrotron Radiation facility (ESRF), Grenoble, France, at wavelengths of 0.8729 and 0.9677Å, respectively. Data indexing, integration and scaling were performed using XDS/XSCALE (Kabsch, 2010). Phases were obtained using ARCIMBOLDO (Rodríguez et al., 2009) (K12A) or by molecular replacement with Phaser (McCoy et al., 2007) (L15A, G16A). For the molecular replacement of G16A and L15A, the structure of WT PSMα3 was used as a search model. Crystallographic refinements were performed with Refmac5 (Murshudov et al., 1997). Model building for all structures was performed with Coot (Emsley and Cowtan, 2004) and illustrated with Chimera (Pettersen et al., 2004). There were no residues detected in the disallowed region of the Ramachandran plot. All PSMα3 mutant structures contained one α-helix in the asymmetric unit. The K12A structure contained three water molecules, and one sulfate anion in the asymmetric unit. The L15A structure contained 11 water molecules and one MRD ((4R)-2-methylpentane-2,4-diol) molecule. The G16A structure contained two water molecules, two sodium cations, and one chloride anion in the asymmetric unit. Two alternative conformations of Leu7 were detected. Crystallographic data and statistics for each structure are listed in Table 2.

### Calculations of structural properties

The inter-sheet distance in the structures was calculated using Chimera (Pettersen et al., 2004). First, a plane was defined for each sheet of α-helices containing at least four helices, using only the backbone atoms of the helix and excluding flanking residues in each structure. Then, the distance between the planes was measured using the “measure distance” command. This method was used to measure distances of both the dry and the wet interfaces. To calculate the distance between α-helices along the sheet, an axis was defined for two adjacent helixes, again using only backbone atoms that participate in the helical portion of the peptide, and the distance between the two axes was measured in the “define axis/plane/centroid” window. All calculated values, as well as the lengths of the axes calculated for the α-helix of each peptide structure, are summarized in Table S1. Of note, the values calculated here for WT PSMα3 are slightly different than those published earlier (Tayeb-Fligelman et al., 2017), due to differences in calculation methods.

### Data availability

The data that support the findings of this study are available on request from the corresponding author. Coordinates and structure factors for the X-ray crystal structures have been deposited in the protein data bank (PDB) with accession codes 6GQ2, 6GQ5, and 6GQC, for K12A, L15A and G16A respectively.

